# Context-dependent mechanical reconfiguration is necessary for multifunctional behavior in a constrained hydrostat

**DOI:** 10.64898/2026.04.01.715937

**Authors:** Michael J. Bennington, Stephen M. Rogers, David M. Neustadter, Roger D. Quinn, Gregory P. Sutton, Hillel J. Chiel, Victoria A. Webster-Wood

## Abstract

Muscular hydrostats, muscular structures with no rigid skeleton, are ubiquitous within the animal kingdom, from vertebrate tongues to cephalopod arms^1,2^, but how they perform complex actions remains poorly understood. One model hydrostat studied for its neural control^3–7^ and biomechanics^8–17^ is the feeding system (buccal mass) of the sea hare *Aplysia* (Fig. 1). The buccal mass (Fig. 1b) performs multiple feeding behaviors by coordinating intrinsic muscles to move a grasper (odontophore)^18,19^. In this paper, we investigated how mechanical reconfiguration from interacting shape-changing elements facilitates large odontophore protractions. During rejection behaviors, mechanical reconfiguration of the odontophore (elongating its shape to a higher aspect ratio) stretches a protractor muscle (I2), allowing I2 to generate stronger protractions^12^. In biting behaviors, the odontophore has a similar range of motion. However, during biting, the odontophore has a lower aspect ratio throughout protraction, meaning the I2 muscle alone is insufficient to reach observed protractions due to its length/tension property and reduced mechanical advantage^9,10,12,18^. By combining new analysis of MRI movies of *Aplysia* feeding^12,18^ (Fig. 1) with a new biomechanical model for biting and rejection (Fig. 2), we demonstrate two context-dependent mechanical reconfiguration mechanisms that explain the different ways large protractions are produced in biting and rejection (Fig. 3). The mechanisms integrate shape changes, bending and conforming of muscle structures, and shifts in contact interactions. We propose two mechanical subclasses of muscular hydrostats, “constrained” or “unconstrained” (Fig. 4), that may be morphologically similar but employ different control strategies depending on whether mechanical constraints are reliably present.

## Results

### Kinematic differences in biting and rejection suggest behavior-specific mechanisms of reconfiguration

*Aplysia* uses its feeding organ, the buccal mass (Fig. 1a-b), in different ways to ingest and reject food. Biting is an ingestive, searching behavior where animals attempt to grab nearby food^19^. To do this successfully, the animal must open the radular surface of its odontophore (the internal grasper of the buccal mass, Fig. 1b) during the protraction phase (Fig. S1), separating the two halves of the surface laterally.

**Fig. 1.**
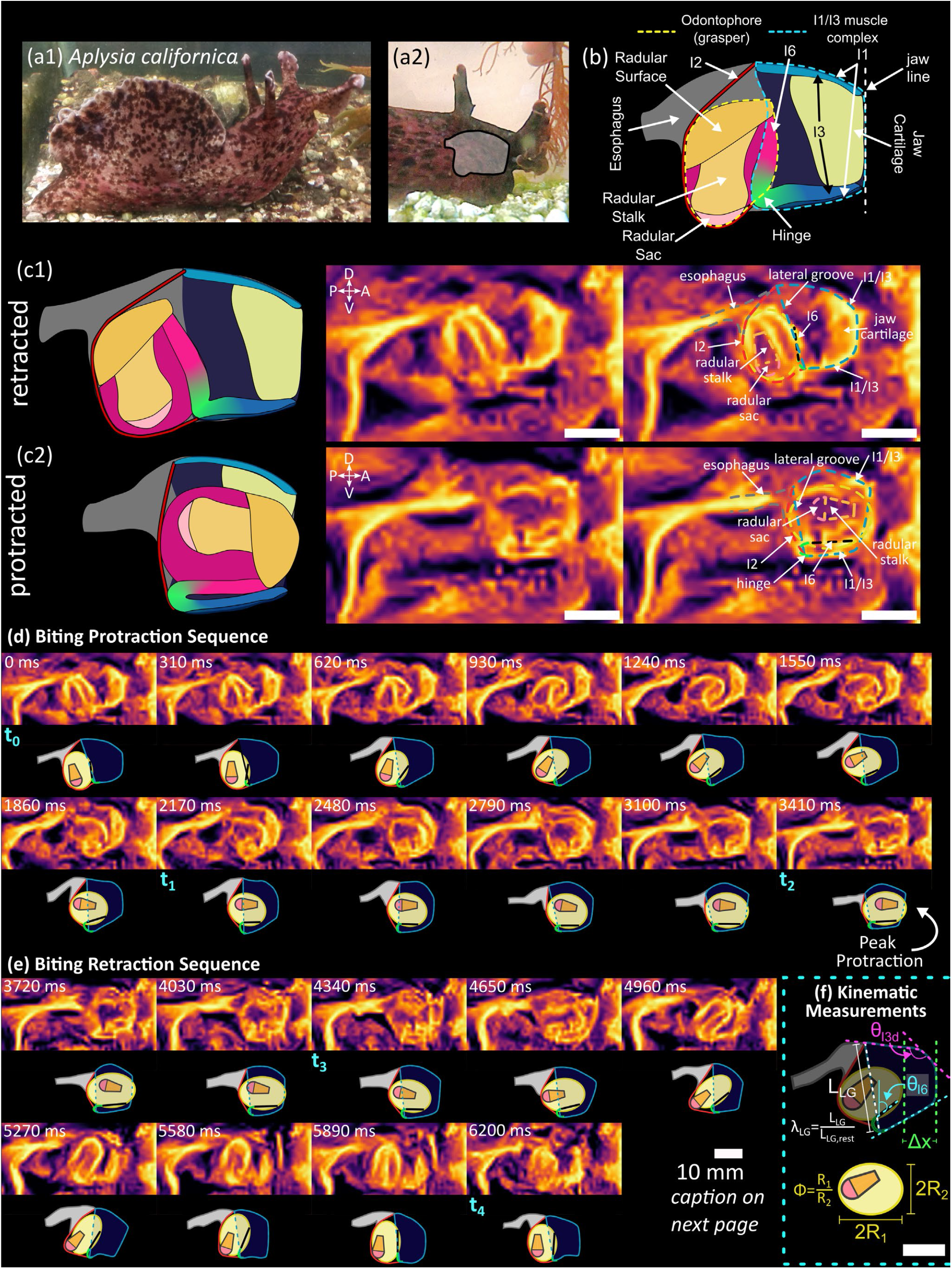
Anatomy and kinematics of the *Aplysia* feeding system. (a1) Adult *Aplysia californica* searching for food and (a2) feeding on *Gracilaria* macroalgae ((a1) photo credit: Dr. Jeffrey P. Gill, (a2) modified with permission from Bennington et al. 2025^14^). Gray highlight shows the location of the feeding structure, the buccal mass (b). (b) An anatomical diagram of a midline sagittal view of a buccal mass. During feeding, the odontophore (the internal grasper of the buccal mass) protracts through the tubelike I3 muscle. In the midsagittal plane, the I3 is visible as two longitudinal elements, but is one continuous structure that runs circumferentially around the buccal mass. The inner wall of the distal I3 is shown in dark blue. The dashed white line shows the jaw line, which is used as the reference for both the translation and rotation measurements. (c) Configuration of the buccal mass (left: anatomical diagram; middle: MRI frames) showing (c1) peak retraction and (c2) peak protraction. (right) A diagram of the buccal mass was created to highlight key anatomical landmarks for each frame of the MRI video showing a complete biting sequence (d-e). The same diagrammatic representations of the landmarks are shown in (d) and (e) for the protraction and retraction portions of the biting sequence, respectively (See STAR Methods). The frames shown in (c1) and (c2) correspond to the 0 ms and 3410 ms frames, respectively, and are the same between the middle and right portions of the figure. Key frames referred to in the text: t_0_: start of the behavioral cycle, t_1_: peak rotation reached, t_2_: peak translation reached, t_3_: rotation plateau ended, t_4_: end of behavioral cycle. (f) Kinematic measurements were taken using the drawn diagrams for each frame in the sequence. See main text for definitions of variables. All scale bars correspond to 10 mm.

This results in the odontophore having a low aspect ratio midsagittally. In contrast, in rejection, previously ingested food is expelled from the buccal mass^19^. To successfully reject food, the animal must close its radular surface on the object being rejected during the protraction phase, resulting in the odontophore having a higher aspect ratio midsagittally. Previous biomechanical analyses have shown that the I2 protractor muscle weakens as the odontophore protracts, due both to its length/tension properties and its mechanical advantage over the odontophore^9,12^. In rejection, this is overcome because the higher aspect ratio of the closed odontophore stretches the I2 muscle more than would an open odontophore^12^.

How does the odontophore (Fig. 1b) achieve large protractions during biting behaviors despite its low aspect ratio and a weakening protractor muscle^9,12^? Do kinematic differences between biting and rejection point to different mechanisms of mechanical reconfiguration in each behavior? To investigate possible kinematic mechanisms of reconfiguration, we produced a new analysis of *in vivo* midsagittal Magnetic Resonance Imaging (MRI) recordings of biting^18^ (Fig. 1, Fig. S2) and rejection^12^ (Fig. S3) behaviors. Each frame of the behavioral sequence was processed in ImageJ (Fiji^20^) to enhance key anatomical features (See STAR Methods), and schematics for each frame were then created by hand-fitting shape primitives and spline curves to the MRI data (Fig. 1c-e, Figs. S2-S3). Finally, kinematic measurements were taken from the schematics for the full behavioral cycle (Fig. 3a). These kinematic measurements (Fig. 1f) include the distance from the anterior edge of the odontophore to the anterior edge of the buccal mass (Δ*x*, referred to as translation), and the rotation of the odontophore, measured as the angle (*θ*_*I6*_) between the I6 muscle and the anterior edge of the buccal mass (referred to as the jaw line). I6 forms the anterior edge of the odontophore, and I3 is the anterior tubelike muscle of the buccal mass through which the odontophore protracts and serves to retract the odontophore (Fig. S1). We also measured the aspect ratio of the odontophore (Φ) and the configuration of the I3 (odontophore retractor) muscle. Specifically, we measured the stretch (*λ*_*LG*_) of the posterior edge of the I3 (anatomically called the lateral groove (Fig. 1c), *λ*_*LG*_=current length / rest length) and the internal angle between the tangent line at the anterior and posterior edges of the dorsal I3 (*θ*_*I3d*_). Key time points in each behavior are identified as ti (biting) and τj (rejection), with the index ranging from 0 to 4 (note that these are different than the indices reported in Neustadter et al. 2007^18^; see STAR Methods).

In both the biting and the rejection behavior, the radular surface of the odontophore underwent similarly large protractions. In biting, the translation range of motion was 0.55 buccal mass lengths (BML), and during rejection, it was 0.53 BML. In both behaviors, the tip of the radular surface translates beyond the anterior edge of the buccal mass. For biting peak protraction beyond the anterior edge was 0.07 BML (t_2_), whereas for rejection it was 0.09 BML (τ_2_).

However, there were differences between the behaviors regarding the aspect ratio, which has important implications for how the protraction was achieved. At peak protraction, the odontophore in biting was more spherical (aspect ratio: 1.29), corresponding to an opened odontophore (Fig. S1). In contrast, the odontophore was more elliptical in rejection (aspect ratio: 1.46), corresponding to a closed odontophore. In addition to extending the long axis of the odontophore itself, this elliptical shape helps to stretch the I2 (odontophore protractor) muscle, enhancing protraction by shifting the I2 muscle higher on its length-tension curve and increasing its mechanical advantage^10,12^. But during biting, the I2 is not sufficiently stretched by the more spherical odontophore to exploit this effect^12^. Thus, another mechanism must be at play.

Further differences were observed in the configuration of the lateral groove. In biting, prior to peak protraction (t_1_ to t_2_), the lateral groove shortened posterior to the odontophore (-45% of its resting length), and θI3d correspondingly decreased (-17°) (Fig. 3a). Together, this suggests that the posterior edge of the I3 was wrapped around the back of the odontophore (Fig 1d). At a similar stage of the cycle in rejection (τ_1_ to τ_2_), this change did not occur (lateral groove shortening: +7%, θ_13d change_: +10°, Fig. 3a). Additionally, in biting, the initial retraction of the odontophore (t2 to t3) corresponded with a re-lengthening of the lateral groove and increasing θ_13d_.

The shortening of the lateral groove and consequent wrapping of the posterior I3 around the rear of the odontophore appears to be a biting-specific kinematic feature, changing how the odontophore and the I3 contact and mechanically interact with each other. We hypothesized that this mechanism will cause protraction of the odontophore, thereby assisting the weaking I2 muscle during biting. However, this mechanism is not seen in rejection, indicating that this reconfiguration may not be necessary for, or may even be detrimental to, rejecting food.

### A hybrid kinetic/kinematic model of the buccal mass reproduces key features of its mechanics

To investigate this hypothesis of behavior-specific reconfiguration, we developed a hybrid kinetic/kinematic biomechanical model of the buccal mass (Fig. 2a). For full details, see STAR Methods. Briefly, in this model, user-controlled kinematic degrees of freedom interact with a quasistatic^14,21^ midsagittal biomechanical model of the buccal mass through elastic contact forces. In the kinetic model, the I2 muscle, whose length-tension properties are modified from Yu et al. 1999^9^, wraps conformally around and applies force to an elliptical odontophore. The odontophore is connected to the rest of the buccal mass by the hinge (the junction between muscles I6 and ventral I3; Fig. 1b (green)), modeled as a beam element^22^. In the kinematic model (Fig. 2c), the aspect ratio of the odontophore (Φ, ranging from 1.1 to 1.8) corresponds to how open (smaller Φ) or closed (larger Φ) the odontophore is. Additionally, the lateral groove stretch (*λ*_*LG*_, ranging from 0.72 to 1.1) sets the shape of the posterior I3. The range of kinematic values was chosen to span the range observed *in vivo* (Fig. 3a). Elastic contact forces from the I3 muscle allow these kinematic subsystems to mechanically interact with the kinetic model. As a first approximation, we did not incorporate the wrapping of the I3 (See Discussion). Anatomical and biomechanical properties of the model were calibrated to existing animal data^11,18^ and models^9^ (Fig. 2b, Fig. S9). To ensure the model would provide meaningful insights into the reconfiguration, we validated its ability to capture key biomechanical and behavioral features of the buccal mass.

**Fig. 2.**
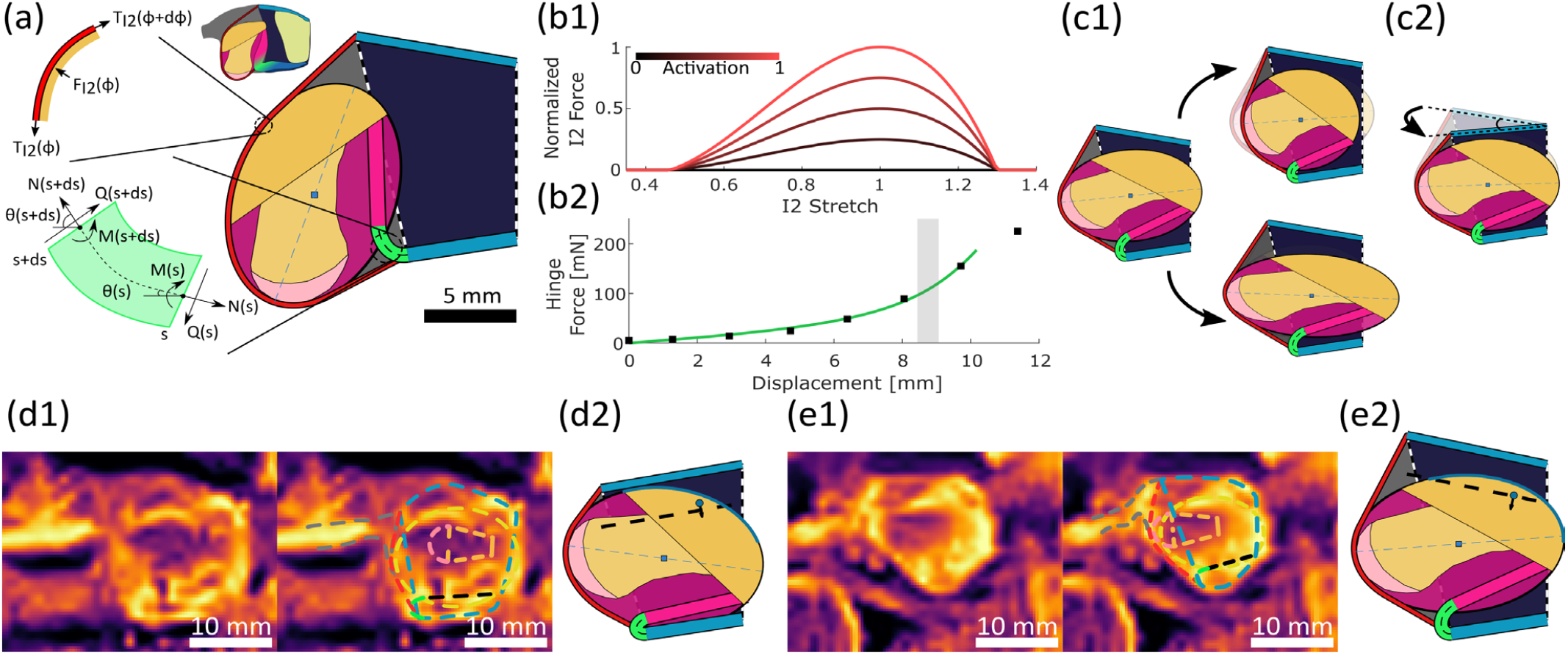
Kinetic/Kinematic biomechanical model of the buccal mass. (a) Rest geometry of the biomechanical model. The grasper (odontophore) is modeled as a rigid ellipse (magenta with yellow radula). It is connected to the I1/I3 lumen (blue trapezoid) by the hinge muscle (green). The I2 protractor muscle (red) wraps conformally around the odontophore and attaches at the lateral groove. The net force and torque from the I2 on the odontophore are found by performing an instantaneous force balance on a small arc of the ellipse and integrating across the full region of contact between the I2 and the odontophore. The hinge muscle is modeled as a linearly elastic, geometrically exact beam. At each position along the beam’s midline, a quasistatic force balance is performed (see STAR Methods). (b1) The tension in the I2 is modeled using the length-tension relationship reported in Yu et al. 1999 scaled by a normalized activation level. (b2) The axial and bending stiffness of the beam hinge were calibrated to *ex vivo* animal data reported in Sutton et al. 2004. Gray region indicates odontophore displacements observed during biting behaviors (Sutton et al. 2004). (c1-c2) To investigate the effects of mechanical reconfiguration on odontophore position at peak protraction, (c1) the aspect ratio of the odontophore ellipse and (c2) the stretch of the lateral groove were added as additional kinematic constraints. (c1) and (c2) show results from the model but do not correspond to any particular behavior or configuration observed in the animal. These constraints impact the biomechanical model via contact forces from the I1/I3 (see STAR Methods). The lateral groove stretch is converted to a depression angle of the dorsal I1/I3 muscle as a proxy for the wrapping of the dorsal I3 around the odontophore observed during *in vivo* feeding behaviors (Fig 1). (d-e) MRI frames at peak protraction in (d1, with and without overlay) biting (t_2_) and (e1, with and without overlay) rejection (τ_2_) compared to corresponding frames from the biomechanical model (d2 and e2, respectively).

We first validated that the model reproduced the hinge’s force response across the odontophore’s full range of motion. Prior models utilizing spring-like muscles either failed to capture the anatomy of the hinge^10,11,13^ or failed to capture the low-displacement force response of the hinge^14^. Re-examination of the hinge’s anatomy (Fig. 1b) led us to hypothesize that the shortcomings were due to a lack of bending stiffness in the modeled hinge^14^. Thus, we modeled the hinge as a beam, combining axial and bending stiffness in a single mechanical element. The beam model of the hinge well approximated *ex vivo* force-displacement responses^11^ (root mean squared error of 3.86 mN across all behaviorally relevant displacements compared to expected force ∼100mN at peak protraction) within behaviorally relevant displacements (Fig. 2b). A sensitivity analysis showed that the hinge forces at low odontophore displacements are due solely to the bending stiffness of the beam, with the axial stiffness contributing only at large displacements (Fig. S4). Together, these results ensure we can predict behaviorally-relevant forces from the hinge and support our hypothesis about the role of bending stiffness in the hinge.

Next, we validated that the buccal mass model could achieve the same peak protraction configurations as observed in the MRI data. By activating the I2 (odontophore protractor) muscle and applying the odontophore aspect ratio (Φ) and lateral groove stretch (*λ*_*LG*_) observed in the MRI data (Fig. 3a), the model successfully reproduces the peak protraction configuration for both biting (Fig. 2d) and rejection (Fig. 2e) behaviors. The model also achieves a similar level of peak translation to the animal data (biting: model peak translation of 0.051 BML, error of 3.8% of the animal’s behavioral range of motion (ROM); rejection: model peak translation of 0.070 BML, error of 4.6% ROM). The model approximates the rotation at peak protraction for biting well (level of rotation: 72.6°, error of 0.76% ROM), but underestimates the rotation in rejection (level of rotation: 64.7°, error of 11.2% ROM). This may be due to deformation observed in the ventral I3 in the MRI data (Fig. 2d-e) that is not modeled with the rigid bar approximation of the ventral I3.

**Fig. 3.**
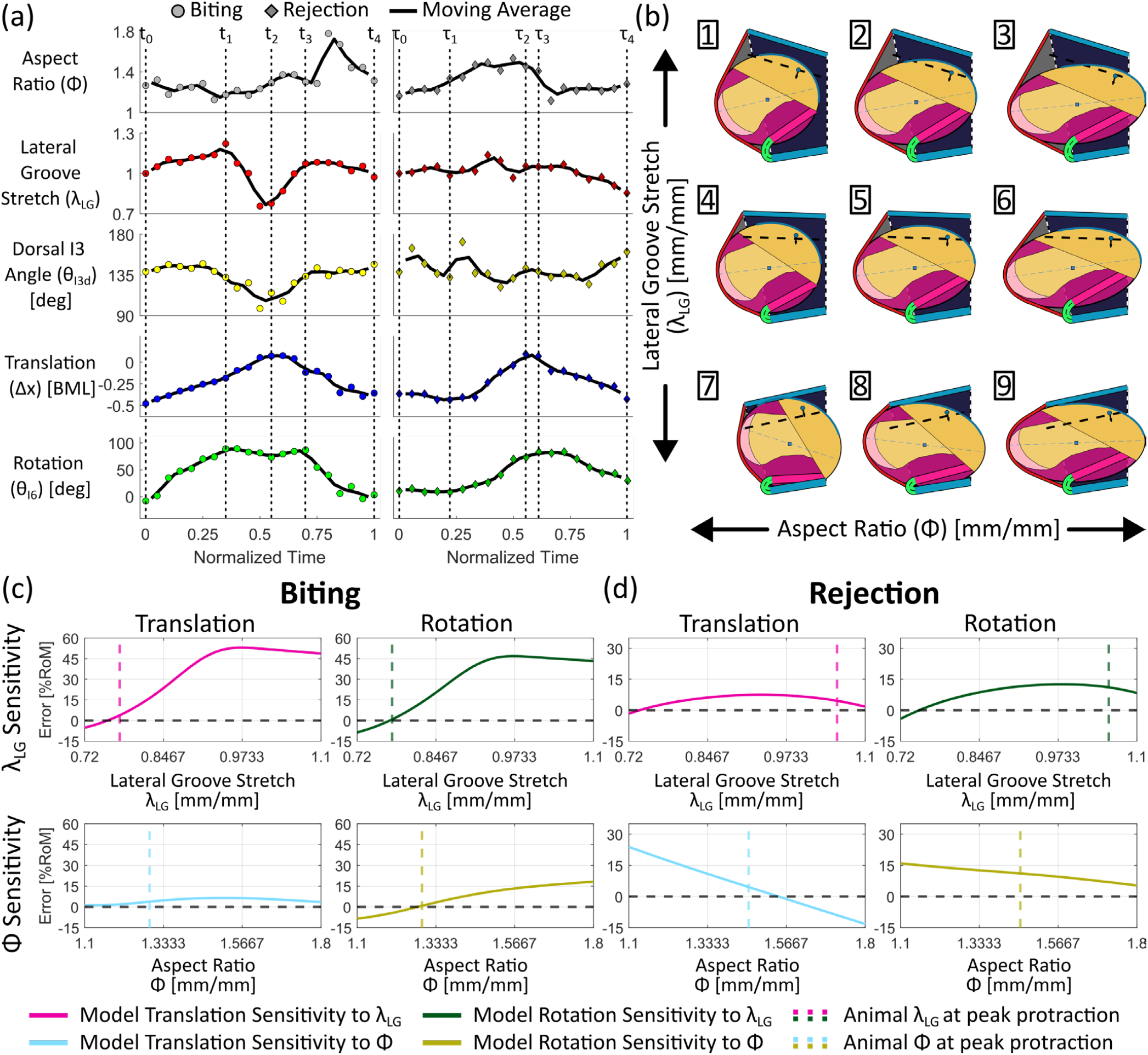
Mechanical reconfiguration of the buccal mass. (a) Midsagittal kinematics of the buccal mass during a (left) biting and (right) rejection behavior (see also Figs. S1 and S2). Colored circles (diamonds) show data for an individual frame, and the black line shows the two-point moving average of the signal. Vertical dashed lines show concurrent time points in the different kinematic signals (biting: t_0_: cycle starts, t_1_: peak rotation, t_2_: peak translation, t_3_: rotation plateau ended, t_4_: cycle ends. Rejection: τ_0_: cycle starts, τ_1_: rotation plateau ends, τ_2_: peak translation, τ_3_: peak rotation, τ_4_: cycle ends). (b) Model configurations for nine different pairs of aspect ratios (*Φ*) and lateral groove stretches (*λ*_*LG*_ ) (numbers correspond to the labeled points in (Fig. S6c)). Note that these simulated results from the model do not necessarily correspond to configurations observed in the animal but rather show changes in the system’s configuration due to changes in the kinematic parameters. All configurations here were achieved with an I2 activation of *A*_*I2*_ = 65%. (c-d) Sensitivity of the model translation and rotation at peak protraction to lateral groove shortening (*λ*_*LG*_, top row) and aspect ratio change (*Φ*, bottom row) for biting (c) and rejection (d). The y-axis for all panels reports the difference between the model prediction and observed animal value at peak protraction (for translation or rotation) normalized by the range of motion (ROM) for each behavior. For each panel, one kinematic parameter is held fixed (top: *Φ* fixed; bottom: *λ*_*LG*_ fixed) at the value observed in the animal at peak protraction, and the other is varied to determine the effect of changing this parameter on the translation and rotation of the odontophore. Vertical dashed lines show the observed value of the varied parameter in the animal at peak protraction. The horizontal dashed line shows 0 difference for reference. The steepness of the difference curve in the vicinity of the vertical dashed line indicates how sensitive the system is to changes in each kinematic parameter near peak protraction. Here, a steeper curve (with a positive or negative slope) indicates greater sensitivity. For biting simulations, *A*_*I2*_ = 15%, and for rejection, *A*_*I2*_ = 90% based on the results of the model validation. Each curve in (c) and (d) is a 1D cross-section of the 2D contour plots shown in Figs. S6-S7. For a complete view of the sensitivity of translation and rotation to lateral groove stretch and aspect ratio across the kinematic configuration space at different I2 activations, see Figs. S6-S7. Note that (c) and (d) use different vertical scales. The smaller scale for the rejection plots was chosen to better show the difference curves for rejection, and it reflects the overall decreased sensitivity to both lateral groove stretch and aspect ratio changes for the rejection behaviors.

Finally, although the level of protraction was similar in both behaviors, the biting configuration in the model could be achieved for a lower activation of the I2 muscle (*A*_*I2*_ ≈ 15%) than was required for rejection (*A*_*I2*_ ≈ 90%) (Fig. S5). A greater I2 activation would take longer to achieve and to relax in the animal, leading to longer overall behaviors. This agrees with and helps to explain observations in animals that rejection behaviors tend to take longer than biting behaviors^23,24^.

### The buccal mass utilizes behavior-dependent reconfiguration to protract the odontophore

From our kinematic analysis of the buccal mass, we observed that the lateral groove shortens during the late protraction phase of the biting behavior (t1 to t2), but not during the rejection behavior (τ1 to τ2) (Fig. 3a). Additionally, the behavior being performed dictates the odontophore’s aspect ratio^12,18^. For biting, the odontophore must be open (lower aspect ratio) during protraction to successfully grab food. In rejection, it must be closed (higher aspect ratio) during protraction to effectively push food out of the buccal mass. Thus, to investigate why lateral groove shortening is used in biting but not rejection, it is necessary to understand how these two mechanisms interact. To investigate the sensitivity of the buccal mass system to these interactions at peak protraction, we performed computational experiments with the model to probe the full kinematic parameter space (Figs. S6-S7). We conducted simulations in which the odontophore was protracted under the action of the I2 muscle, and the model’s kinematic parameters (*λ*_*LG*_ and Φ) were systematically varied while measuring translation (Δ*x*) and rotation (*θ*_*I6*_) of the odontophore. Contour plots of translation and rotation at peak protraction for biting (Fig. S6) and rejection (Fig. S7) across the kinematic parameter space were generated. Finally, cross sections of the contour plots were taken around physiologically relevant configurations to investigate the system’s sensitivity to each parameter at peak protraction (Fig. 3c-d).

When the odontophore is open (as in biting, aspect ratio ∼1.3), the lateral groove shortening leads to odontophore protraction for nearly all levels of I2 activation (Fig. S5-S6). Plotting the difference between the translation achieved by the model for a given lateral groove stretch and the level achieved by the animal at peak protraction reveals the system’s sensitivity to lateral groove shortening (Fig. 3c, top). Specifically, the sensitivity is seen in the steepness of the difference curves near the lateral groove stretch observed in the animal at peak protraction. Conversely, the model is largely insensitive to odontophore shape changes when the lateral groove shortens in biting-like configurations, shown in the much flatter difference curve (Fig. 3c, bottom). Additionally, even for large activations of the I2 (Fig. S6), if the odontophore is open, it cannot reach the translation or rotation observed in the animal without lateral groove shortening. This suggests that with an open odontophore, as in biting behaviors, the buccal mass will require lateral groove shortening to protract the odontophore far enough forward to effectively grab food. Lateral groove shortening is also sufficient to protract the odontophore over a wider range of I2 activations, making it a versatile mechanism for biting behaviors. This supports our hypothesis that lateral groove shortening assists the I2 in protracting the odontophore during biting behaviors. But why is this mechanism only utilized in biting?

When the odontophore is closed (as in rejection, aspect ratio ∼1.5), sensitivity to shortening of the lateral groove at peak protraction decreases. This effect can be seen in the flatness of the difference curves in Fig. 3d (top) and in the contour plot isocurves bending upwards away from the horizontal axis (Fig. S7). The decreasing sensitivity is especially pronounced as the protractor muscle (I2) activation increases. In contrast, the system is more sensitive to changes in aspect ratio (which will change the length of the I2 muscle) (Fig. 3d, bottom). Finally, even at the resting lateral groove length (*λ*_*LG*_ = 1), only I2 activation is needed to achieve large odontophore translations if the odontophore has a high aspect ratio (Fig. S7). This suggests the buccal mass does not need lateral groove shortening to achieve successful protraction of a closed odontophore, as seen in rejection, relying instead on the high aspect ratio of the odontophore to stretch the I2 muscle^12^.

## Discussion

### Bending appears to play a key role in the mechanical reconfiguration of the buccal mass

Previous analyses of the buccal mass and many other biomechanical systems have focused on the stretching of tissue and the bulging of structures^10–14^. However, in this work, we show two key features that rely not on the stretching of tissue along its length, but rather on its bending. In fact, the ability of structures to bend without stretching is critical to their functionality in this system and provides ways for smaller muscles to play an outsized role in the biomechanics of feeding.

The first bending structure is the hinge, connecting the odontophore to the I3, and modeled here as a beam. Previous models of the buccal mass have treated the hinge as an axial spring acting to resist odontophore translation^10–14^. During calibration of this model, we found that at low odontophore displacements, the force from the hinge was only sensitive to the bending stiffness of the hinge (Fig. S4). This implies that, for small displacements, the hinge is only bending and not stretching (the kinematics of the beam in these computational experiments also shows bending alone, Fig. S4). Only at large displacements did the hinge begin to stretch along its length. This bending-dominated kinematics has two effects: 1) the translation and rotation of the odontophore can be coupled in ways not available to purely spring-driven systems, and 2) large translations of the odontophore can be achieved for small input forces without requiring the resisting structure to be weak. Smaller input forces are sufficient because of the mechanical advantage afforded by the moment arm of the load being applied to bend the tissue (Fig. S9).

The second bending structure in this system is the dorsal edge of the I3. For simplicity, we modeled the effect of this wrapping as depression of the dorsal I3. However, in the MRI (Fig. 1), we see that the dorsal I3 conforms to the shape of the odontophore; that is, it bends around the perimeter of the odontophore. This bending and wrapping play key roles in I3’s functionality. First, by wrapping around the back of the odontophore, the posterior contact forces point even more anteriorly, helping to amplify the effectiveness of I3 as a protractor assistant. This would mean that the rigid I3 approximation used in this model may be a conservative estimate of the effectiveness of I3 as a protractor assistant. Second, the wrapping and subsequent contact with the odontophore allow the dorsal surface of the I3 to act as a Class II lever (Fig. S8), with the load acting to protract the odontophore and the effort being supplied by I2. This configuration allows the I3 to serve as a force multiplier for I2 and provides a more advantageous moment arm to further bend the hinge. Traditionally, a lever requires a fixed pivot point (the fulcrum). In the model, the dorsal I3 can pivot about its anterior endpoint (Fig. 2c). However, bending allows for structures to function like a Class II lever but without a true pivot joint, which could not be achieved by a continuous structure like a hydrostat.

The bending of the dorsal I3 may be the result of different interacting components. First, the I2 is well-positioned to apply a bending moment to the I3 as it applies force at a position maximally distant from its base. Additionally, in biting behaviors where this bending is seen, the shape and position of the odontophore means that I2 is positioned more perpendicularly to the average direction of the I3, therefore increasing the resultant moment it generates for any level of activation. Finally, the bending could be influenced by the passive mechanics of the I3 itself and the lateral contact with the odontophore. As the odontophore opens, it will widen mediolaterally, pushing the lateral walls of the I3 outward. This will tend to cause the dorsal and ventral extremes of the I3 to move medially, which in the midsagittal plane would manifest as a bending of the I3. Future studies to investigate these mechanisms will require a 3D odontophore and I3 model and additional coronal kinematic information.

### Lateral groove shortening may be detrimental to the effectiveness of rejection behaviors

The model demonstrated that the buccal mass can protract the odontophore under I2 activation alone when the odontophore is in a closed configuration (as it is in rejection behaviors). Additionally, the shortening of the lateral groove, which is necessary in biting behaviors, is ineffective in protracting the odontophore when the radular surface is closed. So, mechanically, the mechanism isn’t necessary for complete protraction of the odontophore in rejection. From a behavioral perspective, the shortening of the lateral groove and subsequent posterior wrapping of I3 may be disadvantageous for rejection. As the lateral groove shortens, the contact between the odontophore and the I3 increases. This increases the contact force that assists in protraction, but it could also pinch the item being rejected between the odontophore and the posterior I3. This would hold that item in place and make it more difficult for the odontophore to expel it from the buccal mass. This is not a concern in biting behaviors, as biting is a searching behavior performed when there is no food in the buccal mass. Therefore, this mechanism may not be used in rejection both because it is not necessary for the protraction of a closed odontophore and because it may lead to decreased performance in expelling inedible objects.

### Contact interactions guide the kinematics of the buccal mass feeding cycle

From our modeling efforts, we found that the changing contact interactions between the odontophore and the I3 were critical for achieving the correct configurations at peak protraction. Anterior contact between the odontophore and the dorsal lumen of the I3 provided a way for the odontophore to rotate forward and pitch the radular surface down. The changing magnitude and direction of the contact force from I3 also contributed to odontophore protraction for biting-like configurations. Without these contact interactions, the odontophore would not be positioned appropriately at peak protraction to grasp food and successfully feed. Finally, the magnitudes of the contact forces were modulated by other structures (e.g. by the aspect ratio of the odontophore). This modulation allows the contact interactions to play different, context-dependent behaviors.

### Shared mechanical reconfiguration mechanisms suggest there may be subclasses of muscular hydrostats

Muscular hydrostats, with their continually deforming structures, simultaneously offer a remarkable degree of mechanical and behavioral flexibility and present a daunting control task^1,25–27^. Despite this difficulty, animals with muscular hydrostats routinely solve complex control problems to perform flexible behaviors. In other biomechanical systems, the mechanics of the body can offload neural control tasks, thus simplifying the task of descending control^28^. For example, muscle preflexes allow the body to respond to and correct for mechanical perturbations faster than the nervous system can respond^29,30^. The mechanically-mediated stretch activation observed in insect flight muscle^31,32^ allows insect wings to oscillate faster than could be obtained through neural stimulation. Similar stretch activation mechanisms in cardiac tissue allow the heart to modulate its force production to withstand changing preloads and afterloads without direct neural control^33^. Collectively, the role that the body’s mechanics play in its own control is often referred to as morphological computation^34^.

What mechanisms of morphological computation may be at play in muscular hydrostats? The *Aplysia* buccal mass suggests three such mechanisms (Fig. 4a-b). During rejection behaviors, the odontophore actively changes shape by closing the radular surface and increasing its aspect ratio. The resulting stretch of I2 increases the force applied by the odontophore protractor (I2) muscle to the odontophore’s posterior edge, enhancing protraction beyond what neural activation of the I2 alone could produce^12^. During biting, the odontophore changes to its open, more spherical configuration, and the posterior edge of I3 conforms to the odontophore (Fig. 1). The conforming I3 interacts with the odontophore shape change to modulate the contact interaction between the odontophore and the dorsal I3, aiding protraction. Thus, during these two behaviors, the buccal mass uses combinations of 1) actively shape-changing structures, 2) changing mechanical advantage and contact interactions between interacting bodies, and 3) the conformability of structures to other structures.

**Fig. 4.**
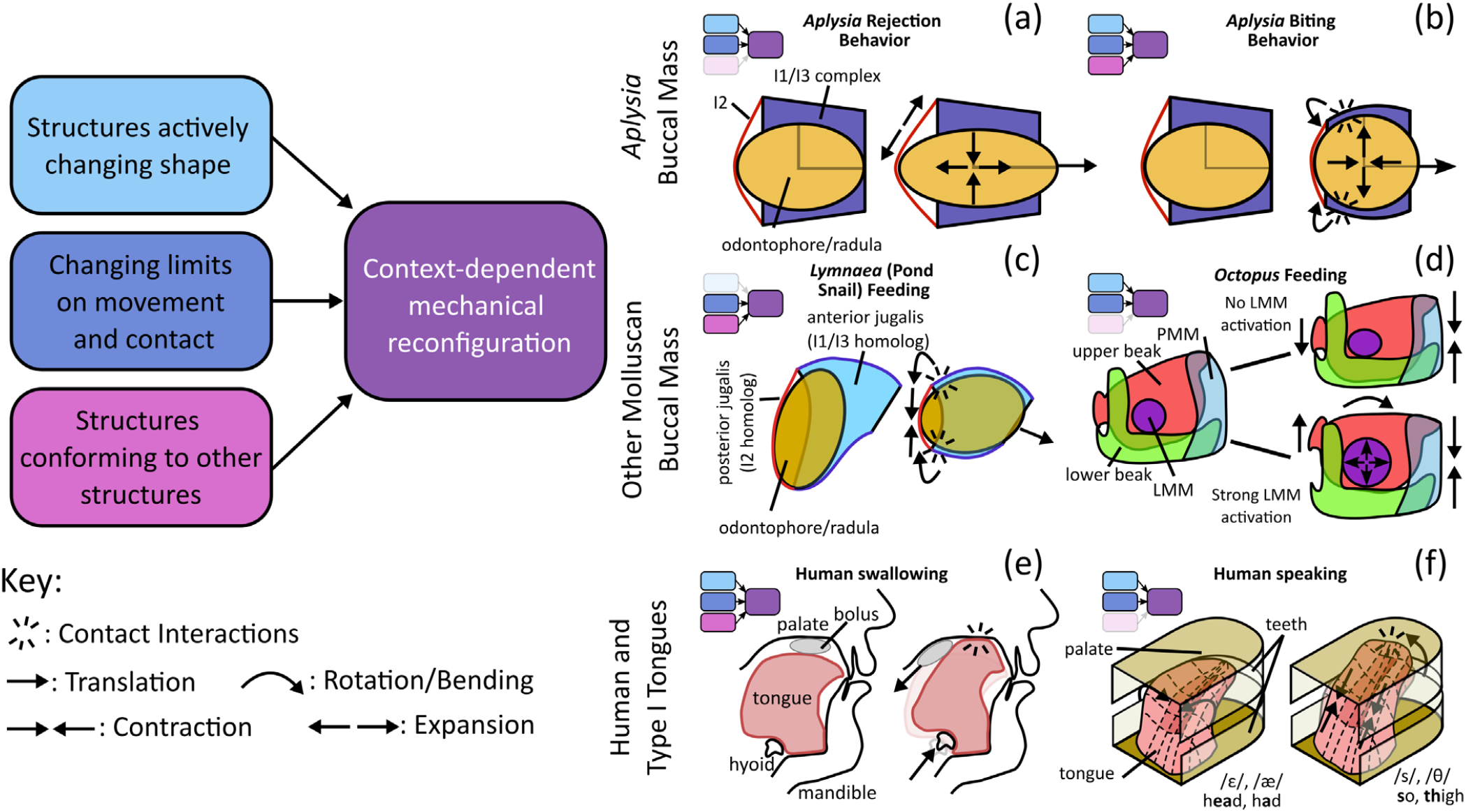
Mechanical reconfiguration facilitates behaviors in a variety of constrained hydrostat systems. Combinations of the active shape change of internal structures (cyan), changes to the movement constraints and contact interaction (blue), and bending and conforming of structures (magenta) allow constrained hydrostats to mechanically reconfigure their neuromusculature (purple) to perform various behaviors. This can be seen in various systems across various species. As discussed here, the *Aplysia* buccal mass uses combinations of these mechanisms in (a) biting and (b) rejection behaviors to protract the buccal mass. (c) The pond snail, *Lymnaea*, has a morphologically similar buccal mass to *Aplysia*, but its I1/I3 homolog, the anterior jugalis, sits further posterior to the odontophore^35^, meaning it may more readily rely on the bending of the anterior jugalis and contact interactions during protraction. (d) The octopus and, more broadly, cephalopod buccal masses contain a beak that lacks a fixed articulation. Instead, by activating the lateral mandibular muscle (LMM), the buccal mass can create a stiff rotation point and may shift the function of the posterior mandibular muscle (PMM) from compressing the buccal mass to opening the beak^36,37^. (e) The human tongue (and other Type I tongues^38^) sits within the skull and makes use of contact with the hard palate to push food from the oral cavity into the pharynx^27,48^. (f) Additionally, by changing how the tongue interacts with the palate and teeth, while maintaining the same internal shape, humans can produce various vowel and consonant sounds^39,49,50^. This use of contact with the palate and teeth is known in the phonetics community as “bracing.” Here, by creating a groove in the middle of the tongue, the phonemes /ε/ and /æ/ can be produced. By raising the tongue and creating palatal contact while maintaining that groove, these vowels shift to the fricative consonants /s/ and /θ/^49^. Small insets show which of the mechanical configurations are used in each behavior.

We believe these mechanisms may be applicable to understanding the mechanics and control of a mechanical subclass of hydrostats we will refer to as “constrained hydrostats” (Fig 4). These constrained hydrostats, e.g., the buccal masses of other mollusks^35–37^ (Fig. 4c-d) and the intra-oral (Type I) tongues of humans and other vertebrates^27,38,39^ (Fig. 4e-f), operate within the confines of other biological structures. These confines limit the range of motion of these hydrostats but also provide them with new opportunities for mechanical interaction (e.g., luminal muscle conformation in *Aplysia* and *Lymnaea*, Fig. 4a-c; muscular articulations^37^ in cephalopods, Fig. 4d; palatal contact in human tongues, Fig. 4e-f). A key feature of the subclass is that, in these hydrostats, the mechanical constraints are consistently present and thus can be reliably used to simplify or constrain the neural control. The odontophore of the *Aplysia* buccal mass always operates within the lumen of the I3; the human tongue predominantly operates within the oral cavity. Because the mechanical interactions (both constraints and opportunities) imposed by the hydrostat’s confines are always present during behavior, the nervous system may encode those interactions, explicitly or implicitly, into the control of the hydrostat’s behaviors. The mechanical environment of constrained hydrostats contrasts with “unconstrained hydrostats,” such as cephalopod arms and elephant trunks. Beyond their attachment to the rest of the body, unconstrained hydrostats are less constricted and predominantly interact with the external environment only. Therefore, they may not be able to take advantage of the same control mechanisms that constrained hydrostats do.

Differences between the mechanical environments of the two subclasses of muscular hydrostats suggest different control strategies. Additionally, hydrostats of the same mechanical subclass may have more similar control mechanisms to each other than to more homologous structures in the opposite subclass. For example, the long and reaching extra-oral (Type II) tongues of, e.g., pangolins^38,40^ may have a more similar control scheme to that of an octopus arm than to the intra-oral control of the human tongue. Interesting model systems to explore the control schemes associated with these two subclasses could be hydrostats that operate both within and without confining structures. These could include the Type I tongues that are also used for food searching and object grasping, like those of cows^38^. These tongues both operate within the oral cavity for food mastication and swallowing, and external to it for grasping and tool use^41^. This subclass hypothesis would suggest that the control schemes used in reaching and grabbing tasks may be more like unconstrained hydrostat control, whereas those used for masticating and swallowing would more closely resemble constrained hydrostat control. More behavioral and computational neuromechanical investigations are needed to thoroughly understand these systems.

### STAR Methods

The detailed methods required to reproduce these studies are available in the accompanying STAR Methods report, including the following sections:

- Key Resources Table
- Resource Availability
  ∘ Lead Contact
  ∘ Materials Availability
  ∘ Data and Code Availability

- Experimental Model and Subject Details
- Method Details
  ∘ Animal Anatomy and Kinematics Imaging
  ∘ Biomechanical Model of the Buccal Mass
  ∘ Parameter Estimation
  ∘ Computational Experiments

## Acknowledgements

This work was supported by NSF DBI2015317 as part of the NSF/CIHR/DFG/FRQ/UKRI-MRC Next Generation Networks for Neuroscience Program. MJB was supported by the NSF Graduate Research Fellowship Program under Grant No. DGE2140739 and the Carnegie Mellon University College of Engineering Presidential Fellowship. Any opinions, findings, conclusions, or recommendations expressed in this material are those of the authors and do not necessarily reflect the views of the National Science Foundation.

## Author Contributions

*Conceptualization*: MJB, SMR, GPS, HJC, VWW. *Data Curation*: MJB, SMR, DMN. *Formal Analysis:* MJB. *Funding Acquisition:* GPS, RDQ, HJC, VWW. *Methodology*: MJB, SMR, DMN, HJC, VWW. *Project Administration:* MJB, VWW. *Resources*: VWW. *Software:* MJB. *Supervision*: HJC, VWW. *Visualization*: MJB, SMR. *Writing – Original Draft:* MJB, VWW. *Writing – Review and Editing*: MJB, SMR, DMN, RDQ, GPS, HJC, VWW.

## Declaration of Generative AI and AI-Assisted Technologies

During the preparation of this work, the authors used Grammarly to improve grammar, spelling, and clarity. No new text was generated using the tools. After using this tool or service, the authors reviewed and edited the content as needed and take full responsibility for the content of the publication.

## STAR Methods Report

**Key Resources Table**

| REAGENT OR RESOURCE | SOURCE | IDENTIFIER |
| --- | --- | --- |
| Biological samples |  |  |
| *Aplysia californica* | Marinus Scientific, Long Beach, CA, USA | N/A |
| Deposited data |  |  |
| Raw and analyzed data (including data generated by the model) | This paper | GitHub Repository (archived: Zenodo) |
| Experimental Models: Organisms/strain |  |  |
| Adult California sea hares (*Aplysia californica*) | Marinus Scientific, Long Beach, CA, USA | N/A |
| Software and algorithms |  |  |
| MATLAB r2024a | MathWorks | MathWorks |
| WebPlotDigitizer v4 | Ankit Rohatgi | WebPlotDigitizer |
| ImageJ Fiji | ImageJ | ImageJ Fiji |
| Inkscape | Inkscape | Inkscape |
| Finite Difference Coefficients Calculator | Cameron Taylor | Finite Difference Coefficient Calculator |

## Resources Availability

### Lead Contact

The lead contact for this work is Dr. Victoria A. Webster-Wood, and all requests for data or further information should be sent to.

### Materials Availability

No new reagents or materials were generated in this work.

### Data and Code Availability

All experimental and computational data generated during these studies, along with the model code and the scripts to conduct all computational studies, are available through [GitHub](http://github.com/CMU-BORG/BuccalMassReconfiguration). An archived version of the code and data is available on <u>Zenodo</u> (doi: 10.5281/zenodo.19373288).

## Experimental Model and Subject Details

All animal kinematic, anatomical, and mechanical data used in this study are from prior experiments (Sutton, *et al*. 2004^11^, Novakovic, *et al*. 2006^12^, Neustadter, *et al*. 2007^18^). One *Aplysia californica* specimen was used to obtain external videos of biting behavior (Fig. S1). This animal was wild caught in Long Beach, CA, USA by Marinus Scientific, and it was kept in a 40-gal artificial seawater (Instant Ocean) aquarium at Carnegie Mellon University (15-16°C, specific gravity 1.023-1.03).

## Method Details

### Animal Anatomy and Kinematics Imaging

To investigate the configurations of the *Aplysia* buccal mass during biting and rejection behaviors, particularly at peak protraction, we analyzed prior MRI video datasets of *in vivo* feeding behaviors. These MRI videos were originally collected as part of the studies reported in Neustadter *et al*. 2007^18^ and Novakovic *et al*. 2006^12^. For complete details of the MRI procedure, we refer the reader to Neustadter *et al*. 2002^42^. Briefly, MRI videos were recorded of the midsagittal plane of the buccal mass during feeding behaviors. Behaviors were elicited using various stimuli, including strips of dried seaweed and noodles flavored with seaweed. All frames took 155 ms to obtain and were separated by 310 ms. One biting sequence (sequence 7521_S4, frames 31-51) and one rejection cycle (sequence 3229_S1, frames 85-103) were chosen to analyze. The biting cycle was chosen as it had not been previously analyzed and a complete feeding cycle could be observed without significant mediolateral motion. The rejection cycle is the same cycle as was investigated in Novakovic *et al*. 2006, as this was the only sequence in the dataset with sufficiently low mediolateral motion or parallax^12^. All MRI frames were post-processed in Image-J Fiji^20^ (Fig. S9). Specifically, the region surrounding the buccal mass was cropped (Fig. S9a), and local contrast enhancement (CLAHE, blocksize=20, histogram=256, maximum slope=3, Fiji^20^) was performed (Fig. S9b). Then the images were upsampled by a factor of 3 using bicubic interpolation^43^ (Fig. S9c) and false-colored using the inverted ‘mpi-inferno’ look-up table (Fig. S9d). For the images used here, we observed that the order of interpolation and contrast enhancement did not make a noticeable difference^43^. The frame showing peak protraction in the biting sequence was then identified as the first frame where the odontophore reaches its furthest level of translation beyond the anterior edge of the buccal mass. Using the MRI frames and the high-resolution lateral view of a bisected buccal mass reported in Neustadter *et al*. 2007^18^, anatomical schematic drawings were created for the rest, retracted, and protracted configurations of the buccal mass using Inkscape. Additionally, for each frame, schematic diagrams were drawn using Inkscape (The Inkscape Project, Boston, MA, USA), depicting the radular stalk, the odontophore outline (drawn as an ellipse), the outline of the I1/I3 complex, the locations of the I6 and hinge muscles connecting to the I1/I3 complex, and the esophagus. Structures were identified in the MRI frames with reference to anatomical images and by comparing sequential frames to differentiate consistent structures from background objects. These MRI frames, in combination with the schematic diagrams, were used for qualitative analysis of the kinematics and to compare with configurations produced by the model. Kinematic measurements were also taken from the schematic diagrams for the full feeding cycle (see main manuscript Fig. 1f). These measurements include:

1. the distance from the front of the odontophore to the jaw line (“translation”, Δ*x*),
2. the angle between the I6 muscle and the anterior edge of the tubelike I3 muscle (“rotation”, *θ*_*I6*_),
3. the length of the lateral groove (and its stretch, *λ*_*LG*_=lateral groove length/lateral groove rest length),
4. the aspect ratio of the odontophore (Φ), and
5. the change in tangent angle between the anterior and posterior edges of the dorsal I3 (*θ*_*I3d*_).

These measurements were made using lines drawn on the schematic diagrams in Inkscape and rescaled from Inkscape units to millimeters using the scale of the MRI images (see Fig. S1 for measurements in biting and Fig. S2 for measurements in rejection).

Key timepoints in the behaviors were then identified from the time series translation and rotation kinematics data (labeled ti (i∈ [0,4]) for biting and τj (j∈ [0,4]) for rejection). The time points are different than the periods identified in Neustadter et al. 2007 (labeled t1 through t4 in that paper). These time points include the start (t0, τ0) and end (t4, τ4) of the cycle, and the time of peak translation (t2, τ2) and peak rotation (t1, τ3). The final identified point is the end of an identified rotation plateau (t3, τ1). For biting, this plateau occurred when the odontophore is protracted, but for rejection, it occurs when the odontophore is retracted. We identified these time points based on the rotation and translation kinematics because these values characterize the gross configuration of the system and are the dependent variables of our model. The other kinematic variables relate to the independent control variables in the model. By identifying key points in the dependent variable time series, we can better identify how these control variables may impact the configuration of the system.

### Anterior View of a Biting Behavior

To visualize the axial view of the animal during feeding behaviors and to show the different configurations of the radula (opened and closed), biting behaviors were elicited and video recorded in an unconstrained animal (Fig. S1). The animal was placed in a behavioral aquarium filled with artificial seawater (16°C). A webcam (Logitech C920, 30 FPS) was mounted on a tripod to provide a top-down view of the behavioral arena. The animal was enticed to the surface of the water with pieces of dried nori. Biting behavior was then elicited by stimulating the lips and anterior tentacles of the animal with pieces of nori attached to wooden dowels. One biting sequence in which the animal faced the camera throughout the behavior was chosen for visualization.

### Biomechanical Model of the Buccal Mass

#### Variable convention

Throughout this document, scalar values are denoted with italicized letters (Latin or Greek), vectors are stylized with an arrow 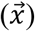, unit vectors are stylized with a hat 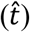, and matrices are indicated with bold typeface (***J***).

#### Mechanics of a beam structure

The “hinge” is the anatomical structure in the buccal mass that connects the odontophore to the I1/I3 complex and is formed by the interdigitation of the I1/I3, I2, I4, and I6 muscles^11^. This interdigitation spans much of the ventral half of the odontophore mediolaterally, and a thick band of tissue runs from the I6 muscle, through the hinge, to the ventral I3^11^. The anatomy of the hinge structure, with its thick midsagittal band and extensive mediolateral components, suggests that this muscle may passively generate forces not only by being stretched but also by being bent or folded. To capture this combined axial and bending stiffness, here we model the hinge muscle as a geometrically exact beam (GEB)^22,44^ (Fig. S10a). A GEB is a 1-dimensional continuum element that can resist both forces and moments and is a geometrically nonlinear extension of classical Euler-Bernoulli Beam Theory. The configuration of the beam is captured by its planar position ((*x, y*)) and its heading angle (*θ*) at each point along the beam (Fig. S10b). In classical formulations and those followed here, a linear constitutive model is assumed, such that axial force and bending moments are linearly proportional to axial strain and curvature, respectively^22,44^. We utilize that assumption here, attributing any observed nonlinearity in the mechanical data to geometric nonlinearities. The governing equations were modified from Voesenek *et al*. 2020^22^ to convert their dynamical equations into quasistatic ones. A quasistatic formulation was used because previous work has shown that *Aplysia* feeding is quasistatic throughout maturation^14,21^. We point readers to Voesenek *et al*. 2020^22^ for a complete derivation of the governing equations.

Briefly, the quasistatic governing equations for the continuum beam element are derived from the instantaneous force and moment balance on each infinitesimal segment of the beam (Fig. S10c). In the absence of body forces (e.g., gravity), this results in the following set of coupled partial differential equations (PDEs).

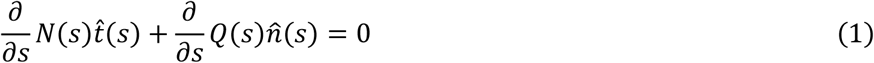

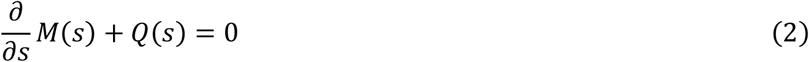

where *s* is the position along the arc length of the beam, *N*(*s*) and *Q*(*s*) are the normal and shear forces at the point *s*, 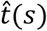 and 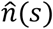 are the tangent and normal vectors of the beam, and *M*(*s*) is the bending moment. These are coupled with constitutive equations

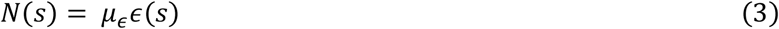

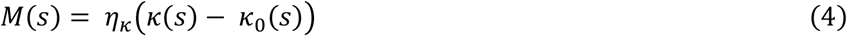

where *μ*_*ϵ*_ and *η*_*κ*_ are the axial stiffness (units mN) and bending stiffness (units mN·mm^2^), *ϵ*(*s*) is the axial strain (unitless), and *κ*(*s*) is the curvature in the beam (units mm^-1^). *κ*_0_(*s*) is the curvature of the beam in the rest configuration. The axial strain is calculated from the position of the beam’s centerline (*x*(*s*), *y*(*s*)) (Fig. S10b) as

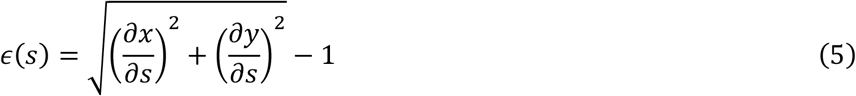

and the curvature is calculated from the heading angle *θ*(*s*) as

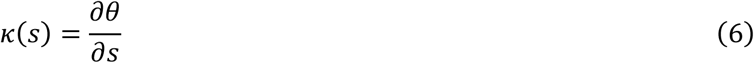

Finally, the heading angle is calculated from the centerline positions as

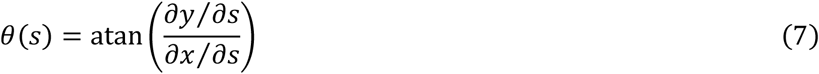

and the tangent and normal vectors can be found using the heading angle as

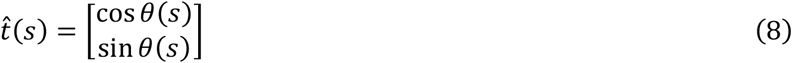

and

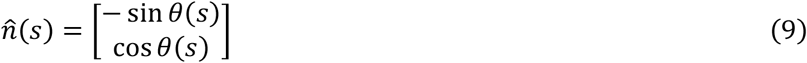

This is a coupled first-order system with the state vector 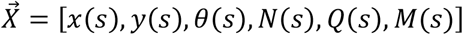. This system requires six boundary conditions to be properly constrained.

#### Hybrid kinetic/kinematic model of the buccal mass

To investigate the effect of buccal mass reconfiguration on grasper protraction in *Aplysia* biting and rejection behaviors, we developed a hybrid kinetic/kinematic model of the buccal mass, where user-specified kinematic degrees of freedom interact with a biomechanical model of the surrounding tissue (Fig. S11). For the work presented here, we assume the buccal mass is bilaterally symmetrical, such that only structures in the midsagittal plane need to be directly modeled, and effects from other structures can be projected onto the midsagittal plane. The buccal mass model consists of a rigid ellipse, modeling the shape of the odontophore, connected to a geometrically exact beam, modeling the hinge muscle structure (Fig. S11a). The tubelike I3 muscle, in the midsagittal plane has a dorsal and ventral element, which we approximate using two rigid bars. The hinge beam connects ventrally to the rigid bar approximating the rest geometry of the ventral I1/I3 complex and dorsally to a rigid bar representing the I6 muscle. The I6 muscle is internal to the odontophore, and, in the model, serves as the connection point of the hinge to the odontophore. Finally, the I2 model is approximated as a chord that attaches to the posterior edges of the dorsal and ventral I1/I3 bars (anatomically called the lateral groove) and wraps conformally around the odontophore.

The forces and torques produced by each model element are used to determine the overall loading on the odontophore ellipse. The tension in the I2 muscle is modeled as a linear combination of a passive tension that is linearly proportional to the stretch 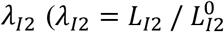, where *L*_*I2*_ and 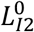 are the current and rest lengths of the I2, respectively) and an active tension. This active tension is modeled using a simplified version of the Hill muscle model presented in Yu *et al*. 1999^9^. Here,

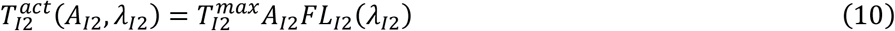

where 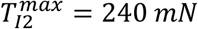 is the maximum isometric force the I2 can generate^10^, and *A*_*I2*_ ∈ [0,1] is the normalized activation of I2. As the dynamics of the system are not considered in this analysis, this activation is simply a control variable that is set during computational experiments. *FL*(*λ*_*I2*_) is the length-tension relationship of the I2 muscle from Yu *et al*. 1999^9^:

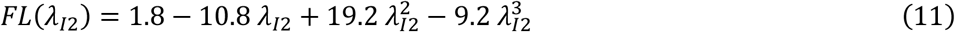

Note that the constant term has been modified from the original paper (originally 1.81) so that the length-tension curve equals 1 at *λ*_*I2*_ = 1. Additionally, in the original Yu model, the length-tension curve was a function of length normalized by the optimal muscle length^9^. Here, we assume the optimal muscle length occurs at its rest length. The total tension I2 is then

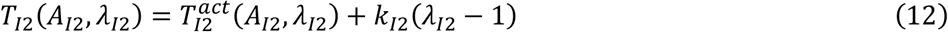

where *k*_*I2*_ = 800 mN is the passive stiffness of the I2 muscle. This value of *k*_*I2*_ was chosen such that the passive force reached 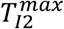 at a stretch of 1.3 as observed in the animal tissue^9^. To translate this I2 tension into its corresponding force vector and torque, we calculate the I2’s mechanical advantage. For a small arc of the I2 where it is in contact with the odontophore (Fig. S7b), the force from the odontophore is balanced by the tension in the I2, which acts tangent to the muscle on each side of the arc. Therefore, the force vector from the odontophore on this arc is equal to the difference in the tangent vectors across the arc. For an infinitesimal arc spanning the elliptical parameter interval *dϕ*, the net force vector on the odontophore, 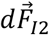, is

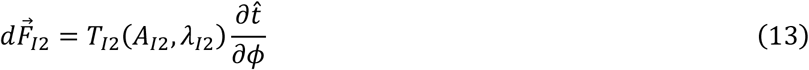

Here, *ϕ* ∈ [0,2*π*] is the parameter of the ellipse boundary, such that for a point (*x, y*) on the boundary of a standard ellipse is found as *x* = *a* cos *ϕ* and *y* = *b* sin *ϕ*. The total force vector that I2 applies to the odontophore is then the integral of 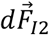 along the arclength of the boundary where I2 is in contact. This simplifies to

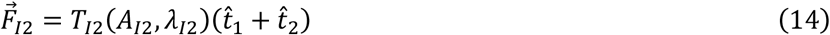

(Fig. S11b). The moment generated by the I2 is

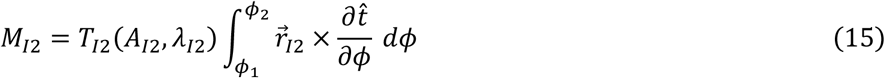

where 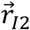 (the moment arm of the arc of I2) is the vector pointing from the hinge attachment point on the odontophore to the point on the odontophore boundary, and *ϕ*_1_ and *ϕ*_2_ are the ellipse parameters where the I2 starts and stops being in contact with the odontophore. Here, × denotes the 2D cross product, from which only the scalar magnitude is considered as the model is planar. This value is integrated numerically (*trapz*, MATLAB 2024a, MathWorks).

The beam hinge model produces passive forces and torques following the constitutive equations outlined above. For the simulations performed here, the active contributions of the hinge were not considered as the hinge is most strongly activated during the retraction phase of behaviors ^11^, and we are focused on protraction-phase configurations of the buccal mass. The beam is parameterized such that the *s* = 0 end connects to the ventral lateral groove, and the *s* = *L* end (where *L* is the arclength of the beam) is connected to the odontophore at the I6 muscle. The boundary conditions for the beam equations are specified using the other components of the buccal mass model. At the *s* = 0 end, the kinematic variables ([*x*(*s*), *y*(*s*), *θ*(*s*)]) are fixed to be equal to their initial conditions (no displacement). At the *s* = *L* end, the kinetic variables ([*N*(*s*) *Q*(*s*), *M*(*s*)]) are specified based on the net forces and moments (about the point where the odontophore connects to the hinge) applied by the other elements of the system to the odontophore. Specifically,

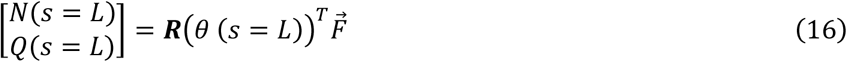

where ***R***(*θ*) is the 2D rotation matrix and 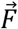 is the total force vector acting on the odontophore.

#### Kinematic constraints and contact forces

To investigate the role that shape changes in the odontophore and the I1/I3 lumen have on protraction, we apply additional kinematic constraints on the system. These constraints are controlled during computational experiments to set different configurations of the system (See Computational Experiments).

The first kinematic constraint relates to the shape of the odontophore and how it attaches to the hinge beam (Fig. S11c). Here, we adopt the odontophore shape change model utilized in Novakovic *et al*. 2006^12^ in which the aspect ratio of the odontophore ellipse (Φ = major axis / minor axis) is the free parameter. The aspect ratio takes values between 1.1 (fully open odontophore) and 1.8 (fully closed odontophore). To determine the lengths of the two axes for a given aspect ratio, we use the same assumptions from Novakovic *et al*. 2006^12^ that the odontophore is an isovolumetric ellipsoid and that the mediolateral radius is the same as the minor axis in the midsagittal plane. Let the major axis be *R*_1_ and the minor axis be *R*_2_. The volume of the ellipsoid is then

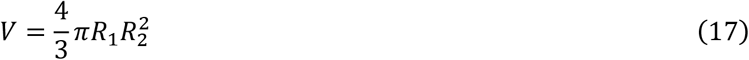

and the aspect ratio is Φ = *R*_1_/*R*_2_. Then, for a specified aspect ratio, we can calculate the major and minor axes as

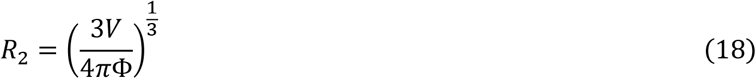

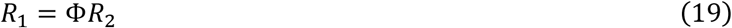

The value for the fixed volume is calculated using *R*_1_ and *R*_2_ in the rest configuration. The location 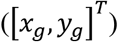 and orientation (*θ*_*g*_) of the ellipse are then determined by the location ([*x*_*H*_, *y*_*H*_ ]^*T*^) and orientation (*θ*_*I6*_) of the *s* = *L* end of the beam hinge (Fig. S11c). We assume that the location of the connection between the beam and the ellipse occurs (1) at the same polar angle from the long axis of the ellipse (*θ*_*H*_: constant) and (2) at a fixed distance from the edge of the ellipse in that same polar direction (Δ*R*: constant) (Fig S11c). Finally, (3) the angle between the I6 bar and the vector from the hinge point and the ellipse center is fixed (*ϕ*_*H*_: constant). These three constraints fully defined the position and orientation of the ellipse for a given configuration of the beam. The orientation of the grasper ellipse (*θ*_*g*_) is found as

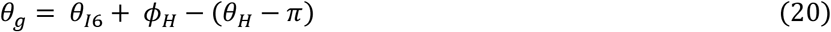

The position of the grasper ellipse is

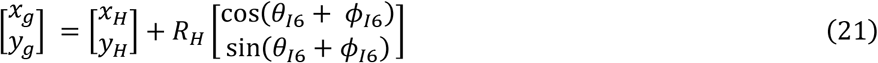

where *R*_*H*_ is the distance from the center of the ellipse to the *s* = *L* tip of the beam:

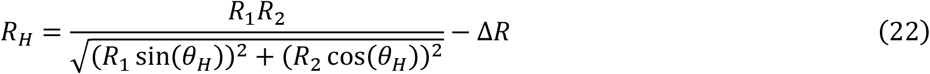

Here, the first term is the effective radius of the ellipse at the polar angle *θ*_*H*_ and will change based on the aspect ratio of the ellipse (through the values of *R*_1_ and *R*_2_). Because this constraint deals only with equality constraints, it is satisfied exactly.

The second kinematic constraint deals with the shape of the I3 lumen and how the I3 contacts the odontophore (Fig. S11d). In the animal, the lateral groove shortens at the peak of protraction during biting^18^. To approximate the effects of this in the model, we posteriorly depress the dorsal line of the I1/I3 complex and simulate the consequent contact forces on the odontophore. Only the dorsal lateral groove point ([*x*_*d*_, *y*_*d*_ ]^*T*^) is changed by this constraint; the jaw location and the ventral I1/I3 line remain fixed. This constraint is applied by specifying the stretch in the lateral groove (*λ*_*LG*_) and calculating the requisite depression angle of the dorsal I1/I3 line relative to the horizontal (Fig. S11d). Specifically, the new dorsal lateral groove point is found as

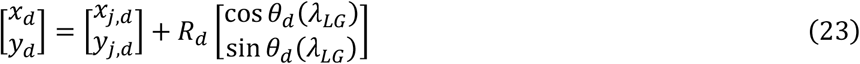

where

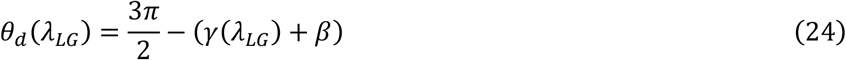

is the angle from the horizontal to the dorsal lateral I3 line, 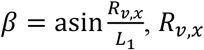, *R* is the x projection of the ventral I3 line, and *L*_1_is the diagonal length of the I3 lumen. Finally,

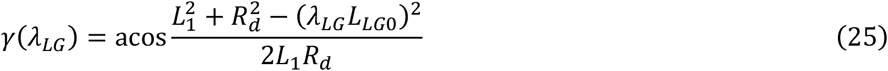

where *L*_*LL*0_ is the rest length of the lateral groove. The effects of this constraint on the odontophore are captured through a contact force (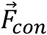, Fig. S11d). In the animal, this contact may not predominantly occur in the midsagittal plane but will be distributed in 3D across the top surface of the odontophore. To approximate this, we assume the contact force is applied to the average position of the ellipse boundary points that are in the top half of the I1/I3 lumen

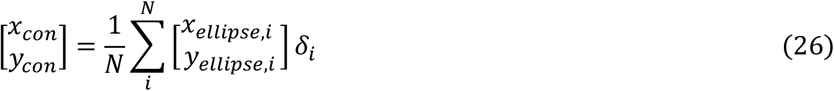

where

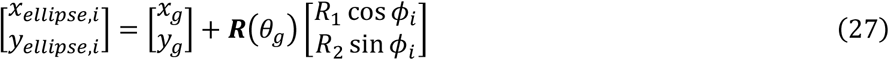

are the boundary points of the grasper ellipse, *N* is the number of points along the grasper ellipse boundary, and

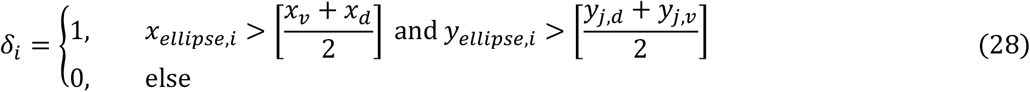

The magnitude of the contact force is linearly proportional to the perpendicular distance from [*x*_*con*_, *y*_*con*_]^*T*^ to a line parallel to and a fixed distance from the dorsal I3 line (white dashed line in Fig. S11d). The contact force always points perpendicularly to the dorsal I3 line.

Finally, to prevent the odontophore from passing through the ventral I1/I3 and to approximate the contact with the ventral I1/I3 complex, a force is applied to any point on the boundary of the ellipse that penetrates the ventral I1/I3 line. The magnitude of the force is linearly proportional to the perpendicular distance of the point to the ventral I1/I3 line. The total force (moment) from this constraint was found by integrating the force (moment) at each point along the boundary.

All constants associated with the kinematic constraints were estimated from the model’s resting configuration using the inverted forms of equations (17)-(25).

#### Implementation of the mechanical model

The governing PDEs of the model were discretized using a finite difference approximation (fourth order approximation for central points, third order approximation for first (last) two boundary points^45^), and the resulting system of equations was simultaneously solved using a custom adaptive implementation of Broyden’s method^46^. Briefly, Broyden’s method is a quasi-Newton method approach to solving nonlinear equations of the form 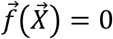, where 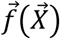 is a system of nonlinear functions of the vector 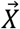. The solution to this system of equations is approximated iteratively using the update rule:

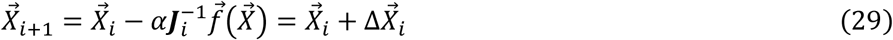

where 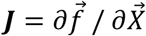 is the Jacobian of the system, and *α* ∈ [0,1] is a scaling factor. Broyden’s method is a quasi-Newton method because, instead of recomputing and inverting the Jacobian of the system at each step, the inverse of the Jacobian is calculated once at the initial condition of the solver and then updated using a rank-1 update based on the step size and change in the function value:

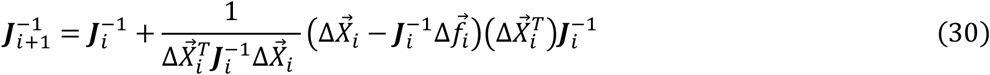

where 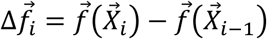 . In the solver used here, the initial Jacobian of the system was also calculated using a finite difference approximation. This solver was made adaptive in two ways. First, within a given timestep of the simulation, to minimize the likelihood of oscillations in the solver, the value of *α* was decreased for each iteration as

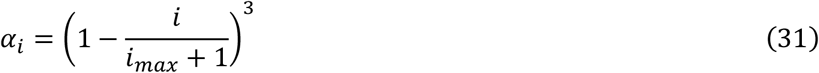

where *i*_*max*_ = 100 is the maximum allowed number of iterations. Second, for a given timestep, if the solver failed to find a solution within the allowed 100 iterations, then the timestep was subdivided into smaller steps, where the number of subdivisions *N*_*sub*_ was iteratively updated as

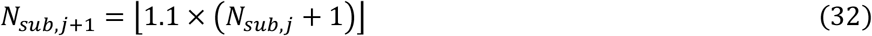

where ⌊*x*⌋ is the floor function.

To avoid numerical instabilities caused by large differences in the order of magnitude of values in the state vector (e.g., between displacements and forces), we adopt the same equation scaling procedure laid out in Voesenek *et al*. 2020 for the PDEs^22^. Based on a convergence analysis, 30 nodes were used to discretize the beam. At this level of resolution, peak force predictions in calibration experiments differed from a reference simulation with 100 nodes by less than 0.1%. All code was implemented using custom MATLAB code (R2024a, MathWorks).

### Parameter Estimation

#### Anatomical parameter estimation

The rest geometry of the biomechanical model was approximated using points and structures digitized from the midsagittal anatomical image presented in Neustadter *et al*. 2007^18^ using WebPlotDigitizer^47^ (Fig. S12a). Specifically, these structures included 1) the boundary profile of the odontophore (magenta squares); 2) the dorsal edge of the I6 (black up-pointing triangle); 3) the anterior edge of the ventral I3 (blue down-pointing triangle); 4) the anterior and posterior edges of the dorsal I3 (blue right-pointing triangles); and 5) a midline curve connecting the I6 through the hinge to the ventral I3 (green circles). The boundary profile points were used to fit the ellipse center and radii (Fig. S11c). To allow the hinge beam to be discretized with any required resolution for the finite difference solver, the midline curve points were fit to a cubic Bezier spline curve that could then be resampled. We parameterized the beam such that *s* = 0 is the ventral edge of the beam connected to the ventral I3 rod, and *s* = *L* is the dorsal edge of the beam connected to the I6 rod.

#### Scale reconciliation

The data used to fit the anatomical and mechanical parameters (See “Constitutive model parameter calibration”) were obtained from animals of different sizes. To account for potential size-dependent effects, we rescaled the anatomy of the model to match the approximate size of the animals used in the mechanical studies presented in Sutton *et al*. 2004^11^. We chose to scale the anatomy and not the force-displacement data because we do not know *a priori* how the force properties of a beam structure will scale with mass, but Rogers *et al*. 2024^21^ recently showed that the midsagittal anatomy of the buccal mass scales isometrically. Data from Rogers *et al*. 2024^21^ were digitized (Fig. S12b) using WebPlotDigitizer^47^ to obtain the appropriate scaling relationships, and the scaling exponents were set for exact isometry (length ∝ mass^1/3^).

To perform this rescaling, the buccal mass masses used in each dataset were estimated, as the animal masses for the individual animals were not reported. From the Neustadter *et al*. 2007^18^ anatomy, we measured both the dorsoventral height and anteroposterior length of the buccal mass according to the measurements reported in Rogers *et al*. 2024^21^ (Fig. S12c). Using the scaling relationships from Rogers *et al*. 2024^21^, we obtained two estimates of the mass of the buccal mass (from height: 4.1 g; from length: 3.2 g), and the mass of the animal’s buccal mass was estimated as the geometric mean of these two values (3.6 g, corresponding to a body mass of 455 g). The geometric mean was used because this would provide the middle point on the logarithmic scale of masses used in the scaling analysis. Then, from the Sutton *et al*. 2004^11^ force-displacement data (See “Constitutive model parameter calibration”), the buccal mass anteroposterior length was obtained from the displacement data, which were reported in both centimeters and buccal mass lengths. Again, using the Rogers *et al*. 2024^21^ scaling laws, the mass of the buccal mass of this animal was estimated as 2.3 g (body mass: 285 g). All length measurements in the anatomical model were then rescaled by (2.3 / 3.6)^1/3^ to obtain the approximate anatomy of the 285 g animal used by Sutton *et al*.^11^. This rescaled anatomy was used for all calibration and computational experiments.

#### Constitutive model parameter calibration

To calibrate the stiffness parameters of the beam model (*μ*_*ϵ*_ and *η*_*κ*_), we simulated the experiments presented by Sutton *et al*. 2004 ^11^. Briefly, in those experiments, the buccal mass was excised, and the I3 lumen was dorsally bisected to expose the odontophore. The odontophore was left connected only by the hinge, and the I3 tissue was pinned out. Then sutures were attached between the odontophore and a force transducer. The transducer was displaced in fixed increments, and the reaction force generated by the hinge was recorded. Displacements were reindexed to the level just after the force was first registered on the force transducer. Data were digitized for the representative animal reported in Sutton *et al*. 2004 Figure 5^11^ using WebPlotDigitizer^47^.

In our simulated experiments, we attached a stiff simulated spring to the dorsal edge of the I6 bar and applied displacements to the other edge of the spring in the direction of the ventral I1 bar. We calculated the displacement of the odontophore as the projection of the displacement of the tip of the ellipse along the direction of the ventral I1 bar. The reaction force was calculated using the stretch of the simulated spring. This allows for the calculation of the analogous force-displacement curve to compare with the Sutton data^11^. The mechanical parameters *μ*_*ϵ*_ and *η*_*κ*_ were optimized to minimize the squared error with the animal data within behaviorally relevant displacements. The squared error was minimized numerically using the MATLAB function *fminsearch* (R2024a, MathWorks).

### Computational Experiments

To investigate the effects of the odontophore and I1/I3 shape change on the protraction of the odontophore, we conducted the following computational experiment using the presented biomechanical model of the buccal mass. Note that the time series of odontophore and lateral groove shape change followed in these experiments do not correspond to a specific feeding behavior but rather allow us to efficiently cover the full kinematic parameter space. As the mechanics of the model and animal system are quasistatic^21^, and therefore not history-dependent, the configurations of the buccal mass obtained in these computational experiments are still relevant to behaviors even if the path between configurations is different. In these experiments, the odontophore is protracted by activating the I2 while the aspect ratio of the odontophore is changed from its rest shape to a specified final aspect ratio. Once the I2 has reached the specified maximum value of activation and the aspect ratio has reached its final value, the length of the lateral groove is changed from 1.1*L*_*LL*0_ to 0.72*L*_*LL*0_, where *L*_*LL*0_ is the rest length of the lateral groove. These extremes were chosen based on *in vivo* kinematics measurements from biting and rejection as to cover the *in vivo* kinematic parameter range. Throughout the experiment, the horizontal distance between the jaw line and the anterior-most point on the odontophore (referred to as the translation of the odontophore, Δ*x*) and the angle between the I6 and the jaw line (called the rotation of the odontophore, *θ*_*I6*_) were recorded as analogous measurements to those reported in Neustadter *et al*. 2007^18^ and those measured during the kinematic analysis in this study. Data were collected during the shortening of the lateral groove for multiple simulations with different final aspect ratios. From these data, we could cover the full behaviorally relevant configuration space of aspect ratio and lateral groove stretch. The 2D translation and rotation data were then interpolated using a local quadratic regression (using the MATLAB *fit* function for a ‘loess’ fit type, span = 0.3, normalize = ‘on’, MATLAB 2024a, MathWorks) to create contour plots of translation and rotation over the configuration space (Figs. S6 and S7). These same interpolated functions were used to create the contour cross-sections reported in the main manuscript (Fig. 3c-d). This process was repeated for maximum I2 activations of 15%, 40%, 65%, and 90%.

