## Supplemental Materials for "Context-dependent mechanical reconfiguration is necessary for multifunctional behavior in a constrained hydrostat"

1 April 2026

### List of Figures:

1. Anterior view of a biting behavior
2. Kinematic measurements for a biting cycle
3. Kinematic measurements for a rejection cycle
4. Relative contributions of bending and stretching to the hinge force
5. Sensitivity of the model protraction and rotation to I2 activation
6. Biting protraction sensitivity to kinematic parameters
7. Rejection protraction sensitivity to kinematic parameters
8. The I2 and I3 form a Class II lever acting on the odontophore
9. MRI image processing steps
10. Mechanics of a geometrically exact beam
11. Model geometry and constraints
12. Estimating model anatomy from midsagittal and scaling data

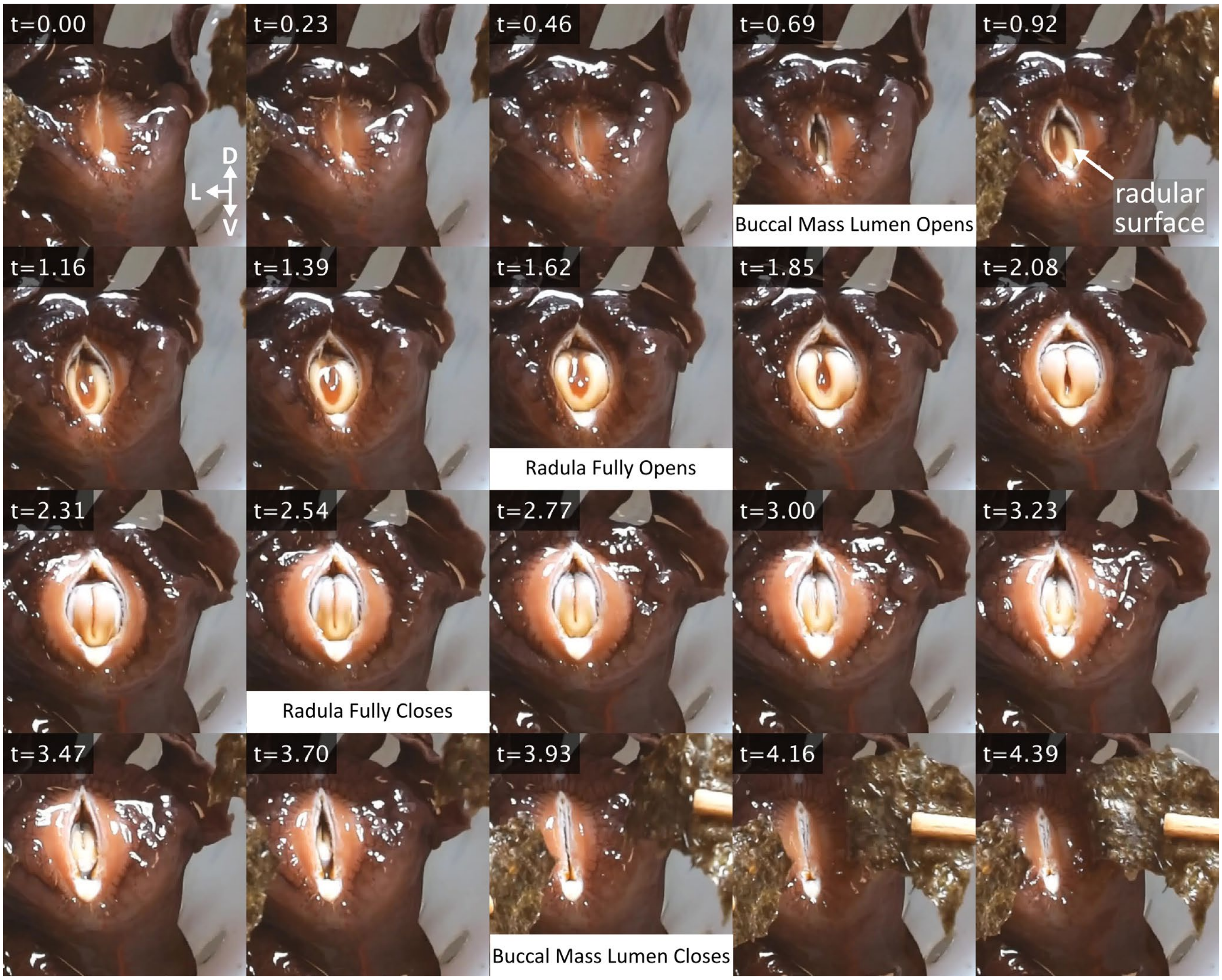


**Fig. S1: Anterior view of a biting behavior** We elicited a biting behavior by stimulating the animal's lips with pieces of dried seaweed. Note, this is a different animal than is reported in the MRI videos (Fig. 1, Fig. S2-S3), so the exact timing and kinematics will vary but are qualitatively similar. At the beginning of the behavior, the jaws and the lumen of the tubelike I3 are closed, hiding the odontophore from external view. When the odontophore has partially protracted, the buccal mass lumen begins to open (t=0.69 s), and the radular surface (tan) can be seen. For biting behaviors, the odontophore protracts with the radular surface open (with its two halves separated laterally). The radular surface reaches its widest extent (~t=1.62 s) just before peak protraction. This open configuration corresponds to a lower aspect ratio shape in the MRI videos. The radular surface then begins to close by bringing the two halves of the surface together medially (t=1.85 to t=2.54 s). Once the radular surface is closed (corresponding to a higher aspect ratio shape in the MRI), the odontophore retracts, and the buccal mass lumen closes. For rejection behaviors, the relative phasing of the radular configuration is reversed, with the odontophore protracting with a closed radular surface (t=2.54 s), opening the radula near peak protraction, and then, once the radula is fully opened (t=1.62 s), the odontophore retracts. Anatomical orientation is shown in the t=0 s frame (D: dorsal, V: ventral, L: lateral).

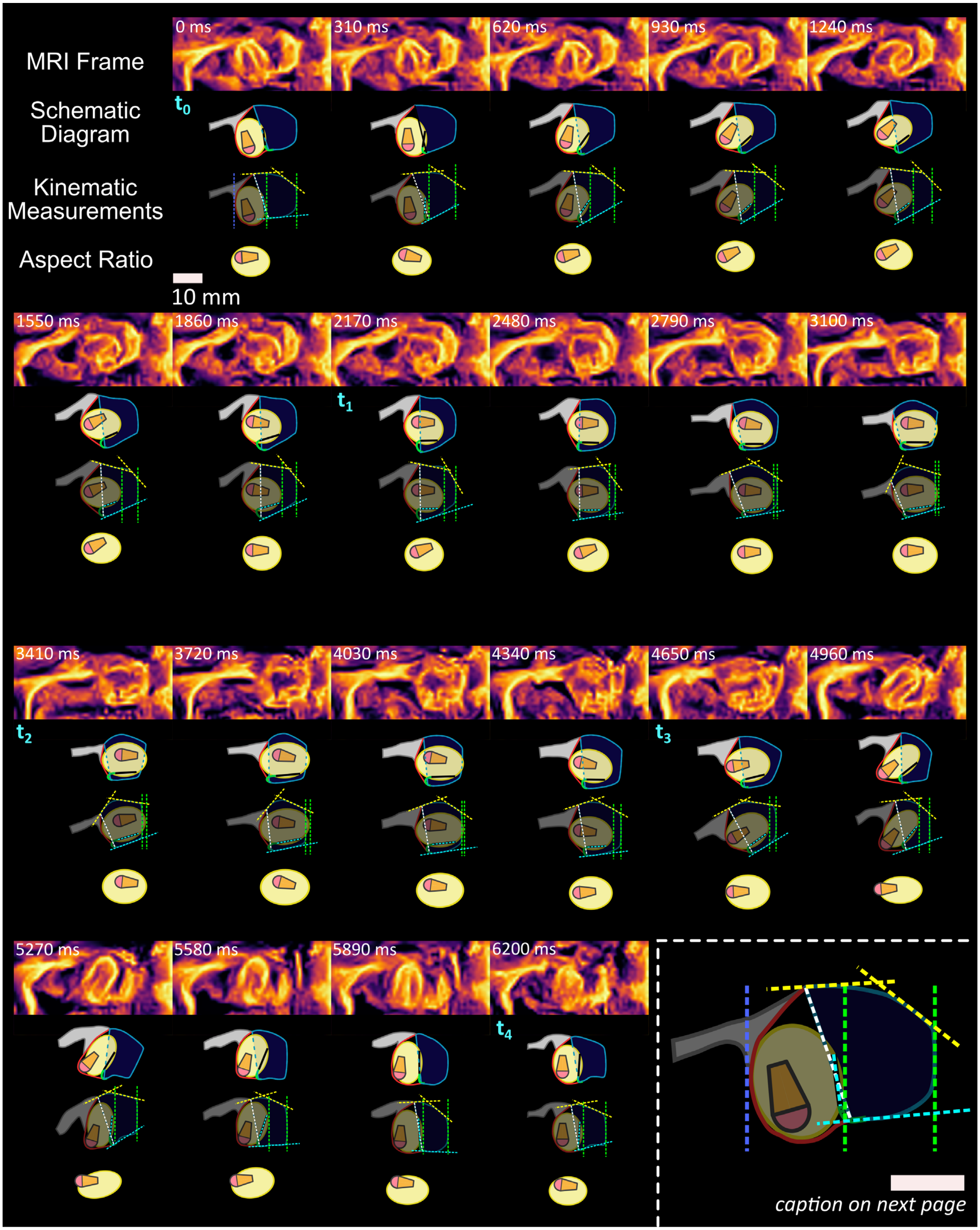

*caption on next page*

**Fig. S2: Kinematic measurements for a biting behavior.** Using MRI recordings of *in vivo* biting behaviors (reported originally by Neustadter *et al.* 2007S1), we measured the midsagittal behavioral kinematics of the buccal mass. The original MRI frames were processed (see STAR Methods and Fig. S4), and schematic diagrams were hand-created using combinations of shape primitives and spline curves in Inkscape. These same frames and diagrams are shown in the main manuscript Fig. 1.

To obtain kinematics measurements from these schematic diagrams, we rotated the schematic diagram so that the anterior edge of the buccal mass was vertical (both to simplify the measurements and to align with the biomechanical model's frame of reference), and a series of lines was drawn on top of the diagrams. (1) Two (green) lines were drawn, one tangent to the anterior edge of the odontophore ellipse and one in line with the anterior edge of the buccal mass; the distance between these lines is the "translation" of the odontophore ($\Delta x$). (2) Two (cyan) lines were drawn, one colinear with the I6 and one aligned with the ventral I3. The angle between the I6 line and the vertical axis (which is parallel to the anterior edge of the buccal mass) is the "rotation" of the odontophore ($\theta_{I6}$). (3) A (white) line was drawn between the dorsal and ventral lateral groove points; the length of this line ($L_G$), divided by the length of this line in the first frame ($L_{G,rest}$), is the "lateral groove stretch" ($\lambda_{LG}$). (4) Two (magenta) lines were drawn, one tangent to the dorsal I3 at the lateral groove and one tangent to the dorsal I3 at the anterior edge of the buccal mass; the internal angle between these two lines is the "dorsal I3 angle" ($\theta_{I3d}$). To measure the aspect ratio of the odontophore (including the radular stalk if it extends beyond the ellipse of the odontophore), a copy of the ellipse and radular stalk was rotated until the width of the object was maximized. In this configuration, the long axis of the odontophore is parallel to the horizontal axis of the page. The "aspect ratio" is then measured as the width of the odontophore ($2R_1$ in Fig. 1) divided by the height of the odontophore ($2R_2$ in Fig. 1). These measurements were repeated for each frame of the MRI sequence. One final (blue) line was drawn in the first frame that is tangent to the posterior edge of the buccal mass; the distance from this line to the green line at the anterior edge of the buccal mass is the "buccal mass length" (BML) for this sample. This value was used to normalize the translation measurement from each frame. An enlarged version (2.5x) of the first frame diagram and lines is shown in the lower right.

These frames were taken from sequence 7521_S4, frames 31-51. This sequence was originally collected during the studies reported in Neustadter *et al.* 2007[S1], but this sequence was not analyzed. The quantitative measurements from this figure are reported in the main manuscript in Fig. 3a. Both scale bars are 10 mm. The smaller scale bar is shared for all frames. The frames labeled with $t_i \in [0,4]$ are the same frames as identified in Fig. 1 and Fig. 3 in the main manuscript, identifying key points in the kinematic cycle (see STAR Methods).

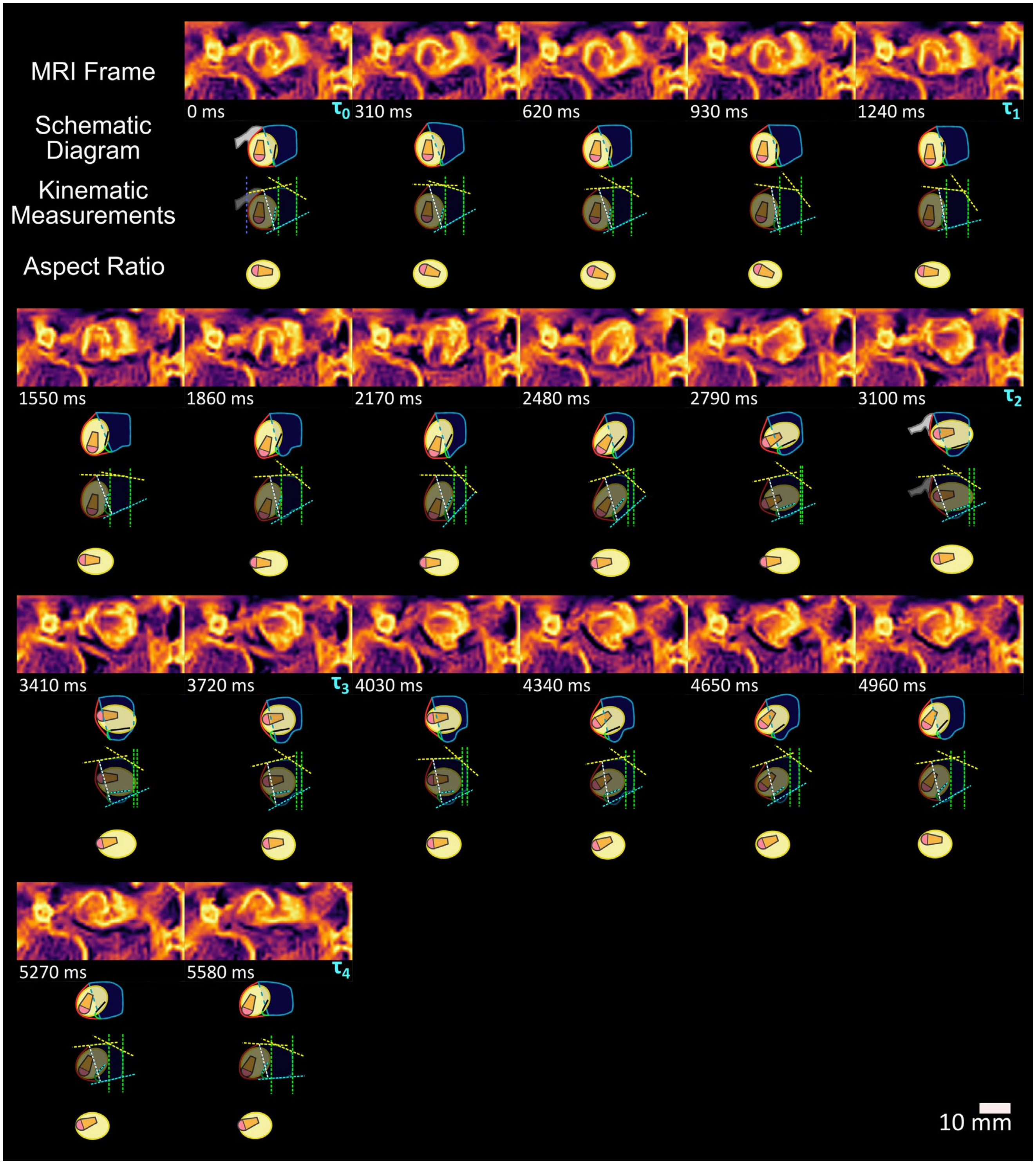


**Fig. S3: Kinematic measurements for a rejection behavior.** The same process used in Fig. S1 was repeated for a rejection behavior. For details of the measurements taken and the associated color coding, see the caption of Fig. S1. These frames were taken from sequence 3229_S1, frames 85-103. This sequence was originally reported in Novakovic *et al.* 2006[S2]. The quantitative measurements from this figure are reported in the main manuscript Fig. 3a. The scale bar is shared for all frames. Frames labeled with $\tau_j \in [0,4]$ correspond to the points labeled in Fig. 3a.

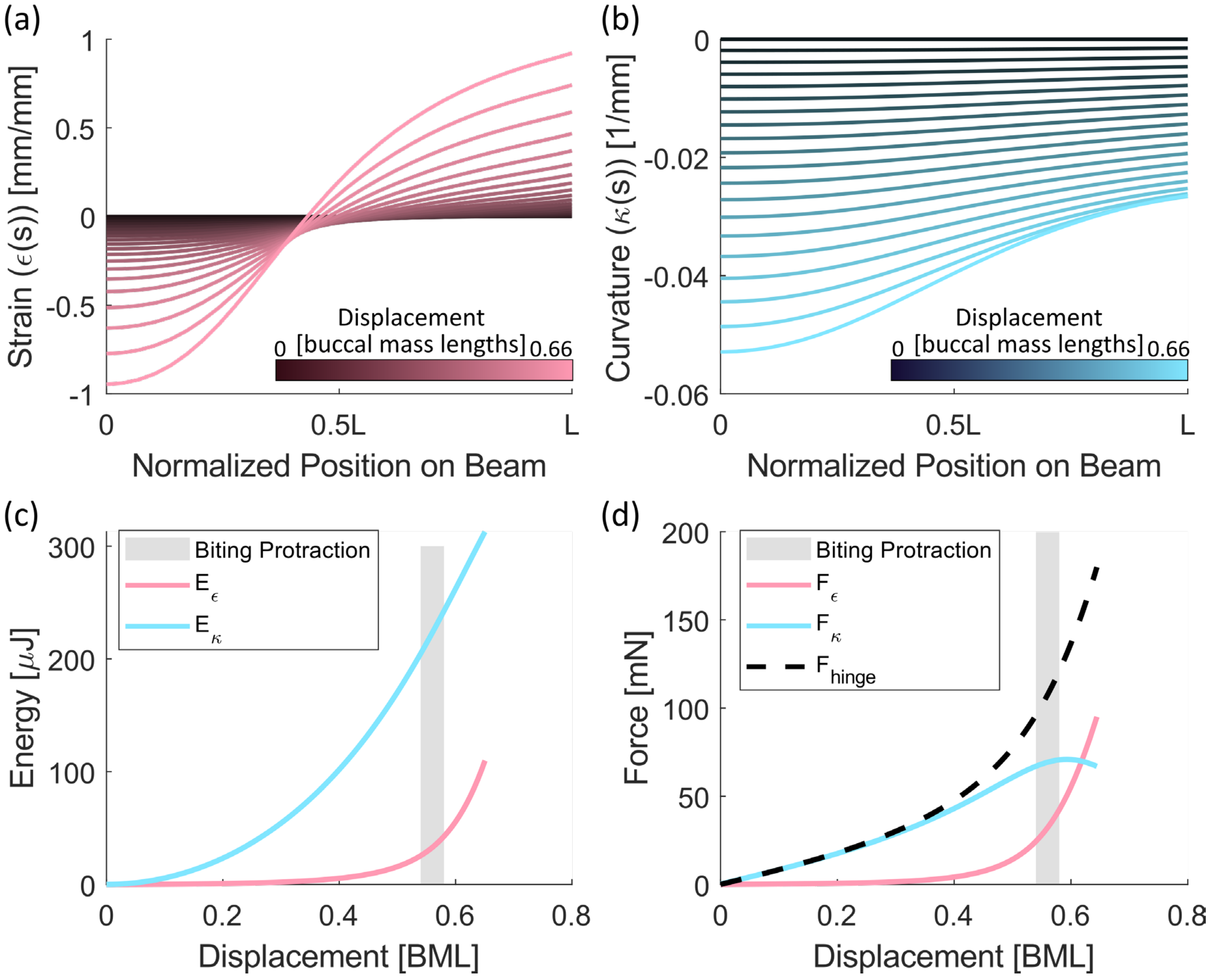


**Fig. S4: Relative contributions of bending and stretching to the hinge force** Using the simulation data from the hinge model calibration experiments (See STAR Methods "Constitutive model parameter calibration"), we determined the relative contributions of bending and stretching to the hinge's force production. For each simulated displacement applied to the buccal mass model, we calculated the distribution of (a) axial strain and (b) curvature along the length of the beam modeling the hinge (beam position is normalized by the length of the beam, L). For small displacements, very little axial strain occurs in the beam, and larger strains only occur for larger displacements. In contrast, the curvature in the beam steadily increases even at low displacements. (c) Using the distributions of axial strain and curvature, we calculated the elastic energy stored in the beam due to axial strain ($E_\epsilon = 0.5\mu_\epsilon \int_0^L (\epsilon(s))^2 ds$) and bending curvature ($E_\kappa = 0.5\eta_\kappa \int_0^L (\kappa(s))^2 ds$). For all displacements of the buccal mass, the elastic energy stored in the hinge is dominated by the bending energy. (d) Finally, we calculated the contributions to the force generated by the hinge on the simulated load cell ($F_{hinge}$) due to the axial strain ($F_\epsilon = \partial E_\epsilon / \partial x_{cell}$) and bending curvature ($F_\kappa = \partial E_\kappa / \partial x_{cell}$). Here, $x_{cell}$ is the position of the simulated load cell along the direction it was displaced during the experiments. Within behaviorally relevant displacement (gray region indicates protraction levels expected during biting behaviors), the force from the hinge comes predominantly from the bending of the hinge. This is especially true for low displacements of the grasper. BML: buccal mass lengths.

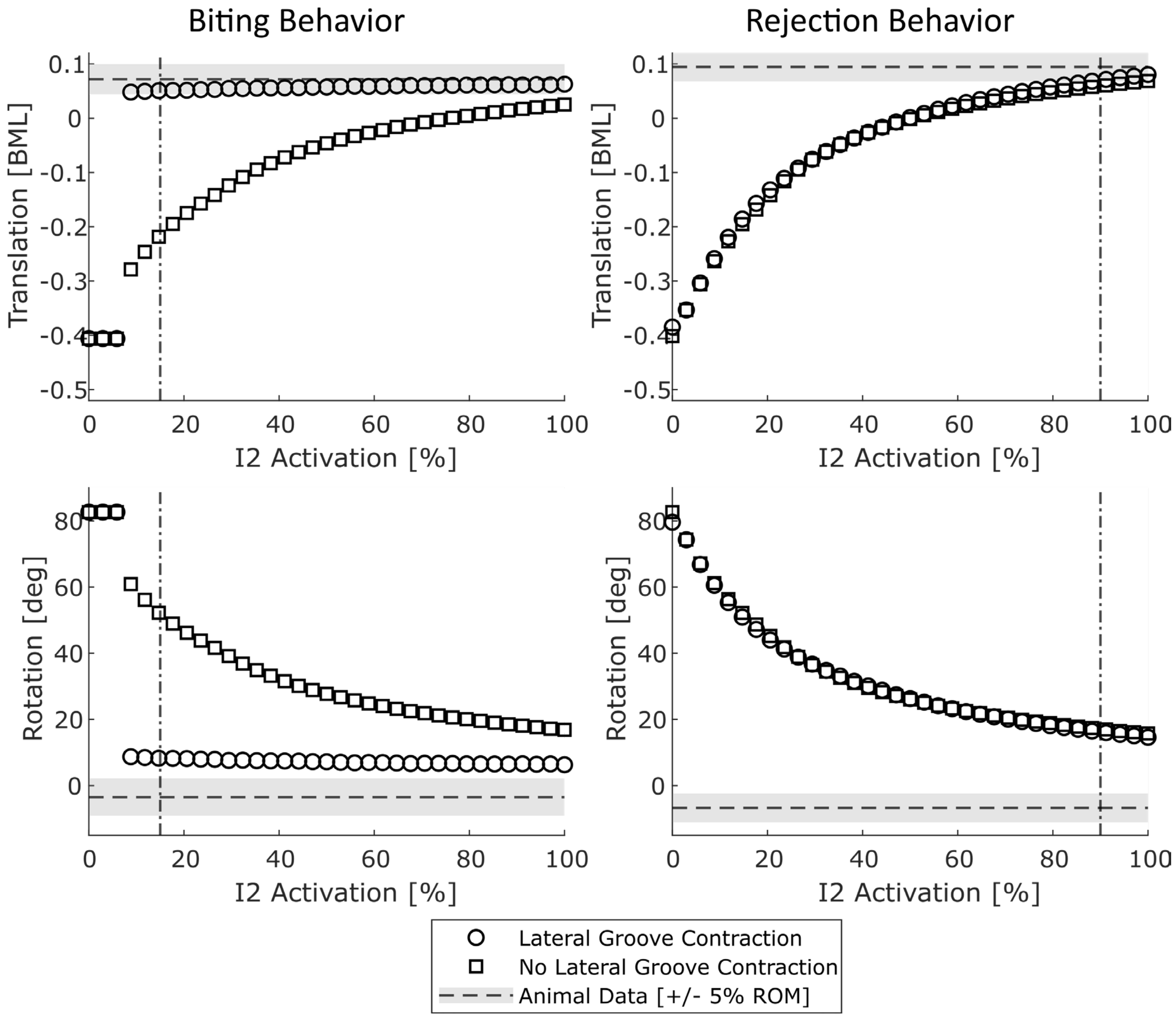


**Fig. S5: Sensitivity of the model protraction and rotation to I2 activation** For biting-like (left) and rejection-like (right) configurations of the grasper, the translation (top) and rotation (bottom) of the odontophore were measured in simulation for prescribed activations of the I2 protraction muscle both with (circles) and without (squares) lateral groove contraction. The aspect ratio and magnitude of lateral groove contraction for each behavior were set at the measured *in vivo* level. The dashed lines show the level of translation or rotation achieved by the animal, and the shaded regions show a range of ±5% of the behavioral range of motion for that metric and behavior. The dot-dash lines show the level of I2 activation used in the simulations reported in the main manuscript Figs. 2d and 2e. In rejection behaviors, the introduction of the lateral groove contraction has minimal effect on the kinematics of the odontophore, and most of the protraction relies on the I2 activation. On the contrary, in biting behaviors, above a critical value of ~9% activation (enough activation to push the odontophore partially into the I3 lumen), the contraction of the lateral groove is sufficient to protract the odontophore the rest of the way to near the *in vivo* level for any level of I2 activation. BML: buccal mass length.

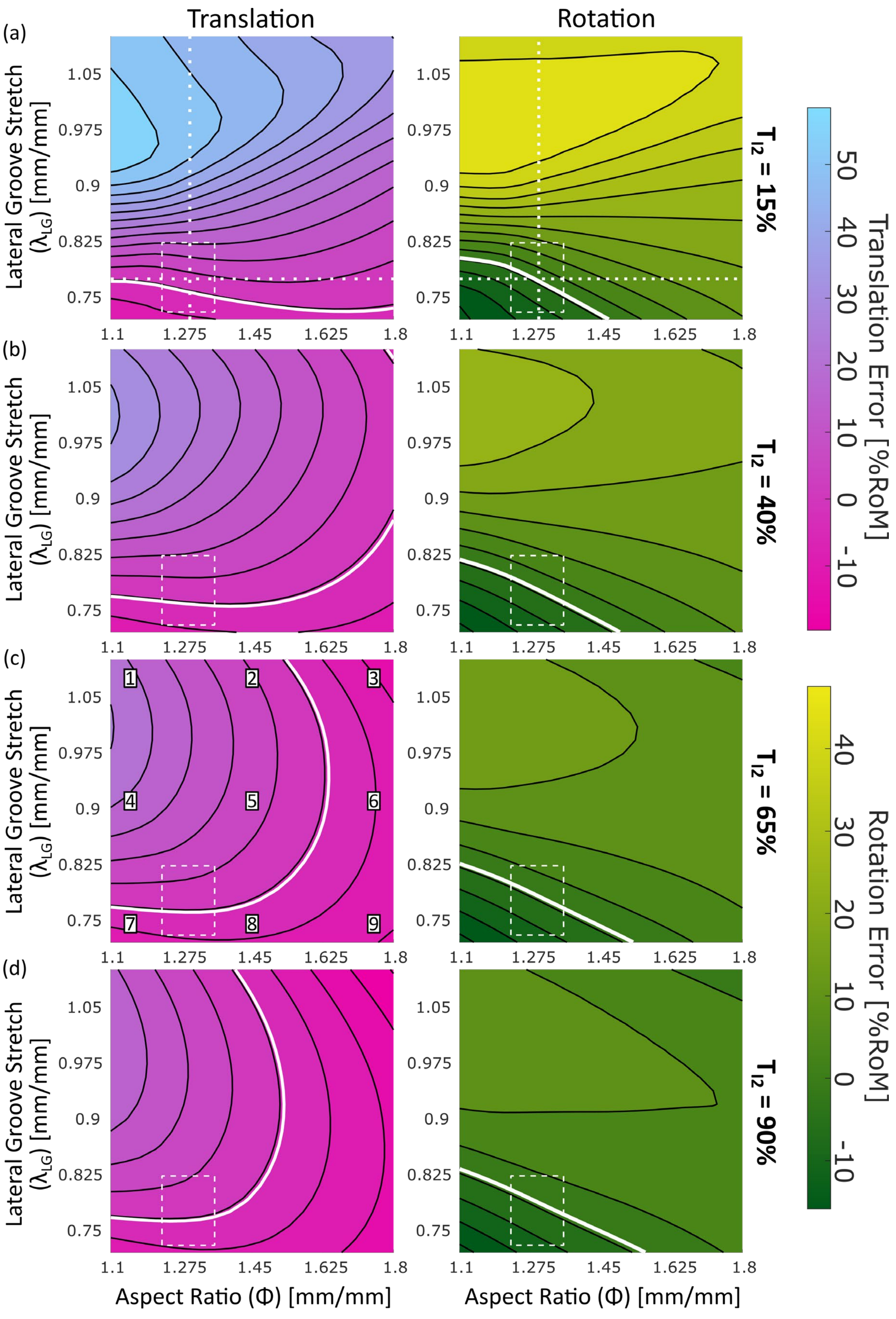

Translation
Rotation
(a)
(b)
(c)
(d)
Lateral Groove Stretch ($\lambda_{LG}$) [mm/mm]
1.05
0.975
0.9
0.825
0.75
1.1
1.275
1.45
1.625
1.8
Aspect Ratio (Φ) [mm/mm]
$T_{I2}$ = 15%
$T_{I2}$ = 40%
$T_{I2}$ = 65%
$T_{I2}$ = 90%
Translation Error [%RoM]
50
40
30
20
10
0
-10
Rotation Error [%RoM]
40
30
20
10
0
-10
1
2
3
4
5
6
7
8
9

**Fig. S6: Biting protraction sensitivity to kinematic parameters.** The simulated levels of odontophore translation (left) and rotation (right) at peak protraction were determined for different combinations of the kinematic parameters $\lambda_{LG}$ (lateral groove stretch) and $\Phi$ (odontophore aspect ratio). These simulations were conducted with $T_{I2}$ = (a) 15% (estimate for biting activation), (b) 40%, (c) 65%, and (d) 90% (estimate for rejection activation). The levels of translation and rotation are reported as the difference relative to the value observed in the animal at peak protraction during a bite (Fig. S2) and normalized by the behavioral range of motion (ROM). Isocurves are equally spaced with a separation of 5% ROM. The color scales for translation and rotation, respectively, are the same across all levels of I2 activation. The white isocurve corresponds to the animal's level of translation/rotation. The white dashed box shows the region of the $\Phi \times \lambda_{LG}$ configuration space near the observed configuration at peak protraction in the animal. The box is centered on the kinematic parameters observed in the animal at peak protraction, and the width and height span ±10% of that parameter's range of motion that was observed during a biting behavior. The white dotted lines in (a) correspond to the cross-sections plotted in Fig. 3c of the main manuscript. Finally, the numbers in (c) show the location in configuration space of the model frames labeled with the corresponding numbers in Fig. 3b of the main manuscript.

From these simulations, we can determine how altering the kinematic variables leads to different levels of translation and rotation, and how sensitive the system is to changes in the kinematic parameters. On the contour plots, areas where the contour lines are primarily horizontal indicate minimal sensitivity to changes in aspect ratio, whereas regions where the contour lines are primarily vertical show minimal sensitivity to changes in lateral groove stretch.

**Fig. S7: Rejection protraction sensitivity to kinematic parameters** (next page). This figure shows the same simulations as in Fig. S6, but with the differences calculated and scaled based on a rejection behavior. The black and white isocurves and the white dashed box show the corresponding information from Fig. S6 (see that caption for details). The white dotted lines in (d) correspond to the cross-sections reported in Fig. 3d of the main manuscript. Note that, while the white isocurves in (d) do not pass through the configuration observed in the animal (the center of the white box), the difference between the model and animal data for translation is still <5% ROM. The difference is 11.2% ROM for rejection rotation, with the higher error likely due to the unmodeled shape change of the ventral I3 that is observed in the MRI (Fig. S3). On the contour plots, areas where the contour lines are primarily horizontal indicate minimal sensitivity to changes in aspect ratio whereas regions where the contour lines are primarily vertical show minimal sensitivity to changes in lateral groove stretch.

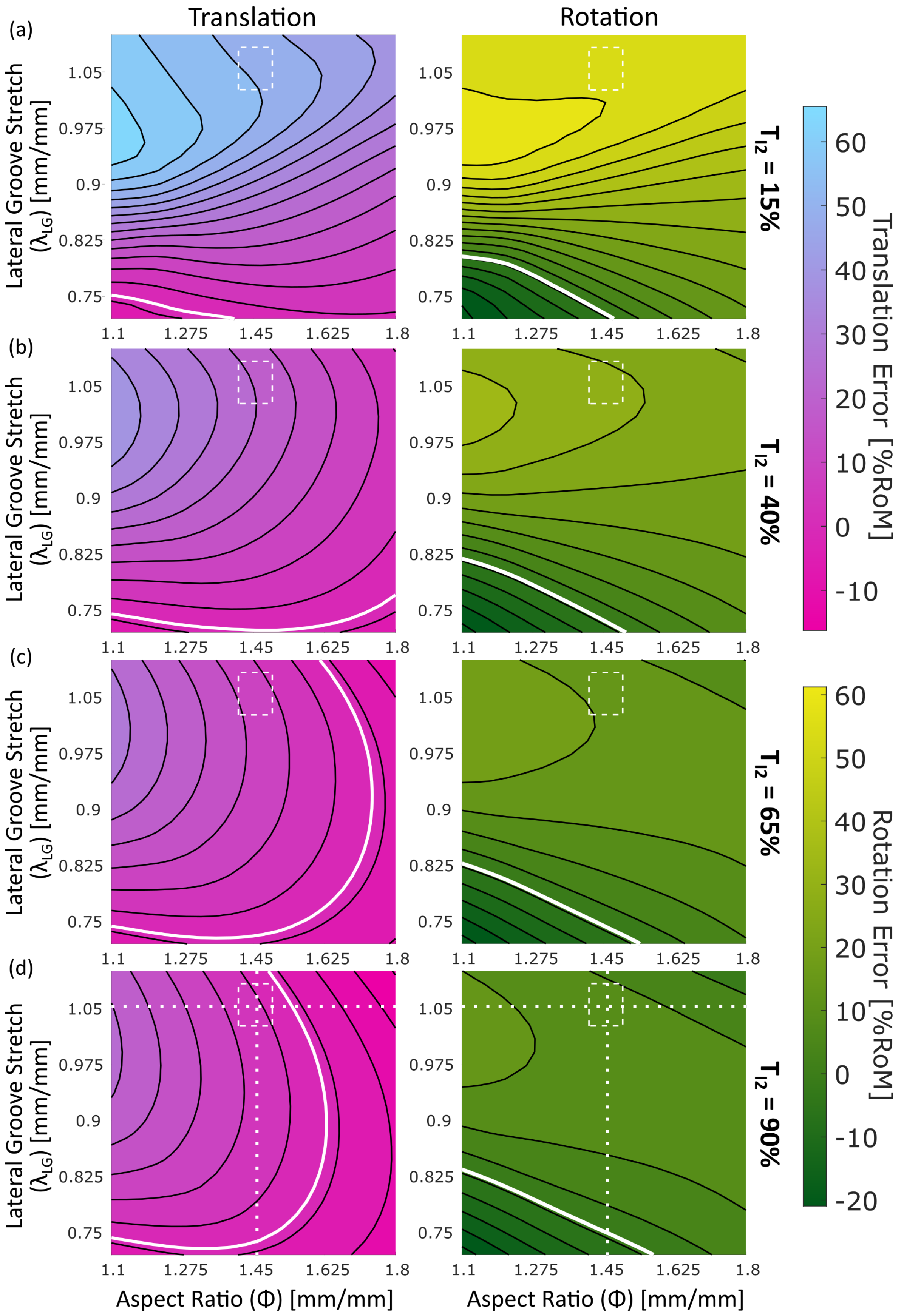

Translation
Rotation
(a)
(b)
(c)
(d)
Lateral Groove Stretch
(λLG) [mm/mm]
1.05
0.975
0.9
0.825
0.75
1.1
1.275
1.45
1.625
1.8
Aspect Ratio (Φ) [mm/mm]
Tl2 = 15%
Tl2 = 40%
Tl2 = 65%
Tl2 = 90%
Translation Error [%RoM]
60
50
40
30
20
10
0
-10
Rotation Error [%RoM]
60
50
40
30
20
10
0
-10
-20

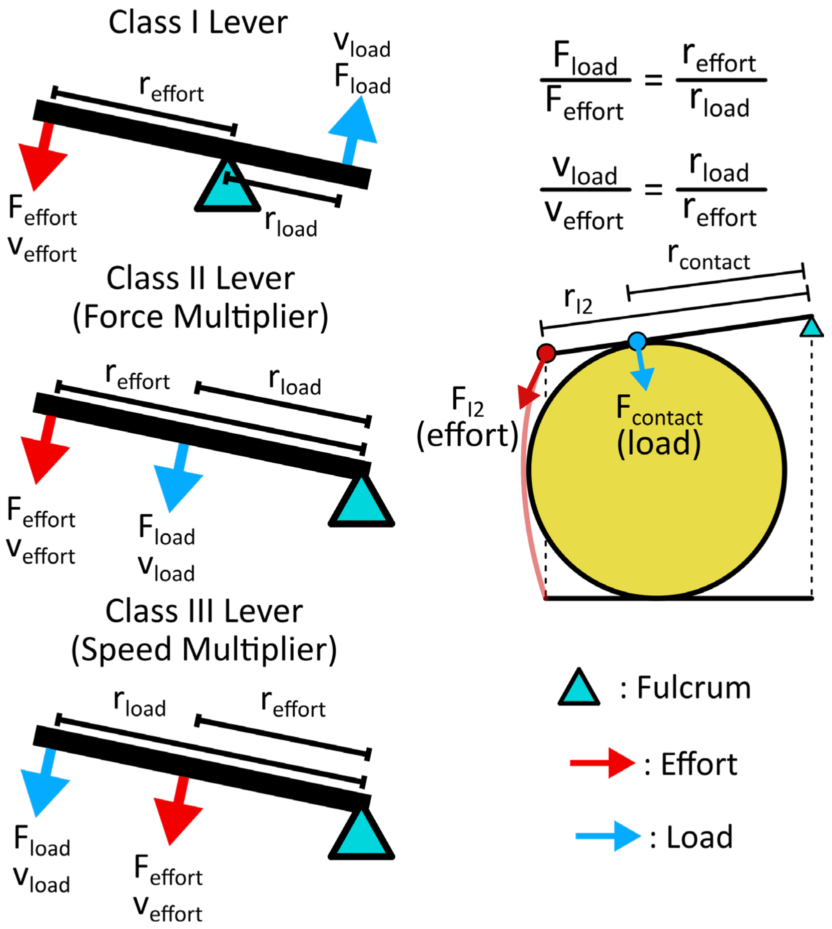


**Fig. S8: The I2 and I3 form a Class II lever acting on the odontophore** By changing the location of the fulcrum (the point about which the lever rotates), the effort (the force applied to do work), and the load (the force against which the effort is working), different classes of levers can be created (left)[S3]. Class I: effort and load are on opposite sides, and whether force or speed is multiplied depends on the ratio of the moment arms (r). Examples include seesaws and scissors. Class II: effort and load are on the same side, with the effort having a greater moment arm ($r_{effort} > r_{load}$), and the load force is always greater than the effort force (force multiplier). Examples include wheelbarrows. Class III: effort and load are on the same side, with the load having the greater moment arm ($r_{load} > r_{effort}$), and the load velocity is always greater than the effort velocity (velocity multiplier). Examples include tongs and human elbows. Right: simplified schematic of the odontophore (yellow circle) contacting the I3 muscle (black lines), with the I2 (red line) pulling on the I3. The I2 muscle applies an effort force to the dorsal I3 muscle. The contact force between the I3 and the odontophore serves as the load. Because the load force has a smaller moment arm than the effort force, the I2 and I3 act as a Class II lever. This amplifies the force coming from I2 acting on the odontophore ($F_{contact} > F_{I2}$).

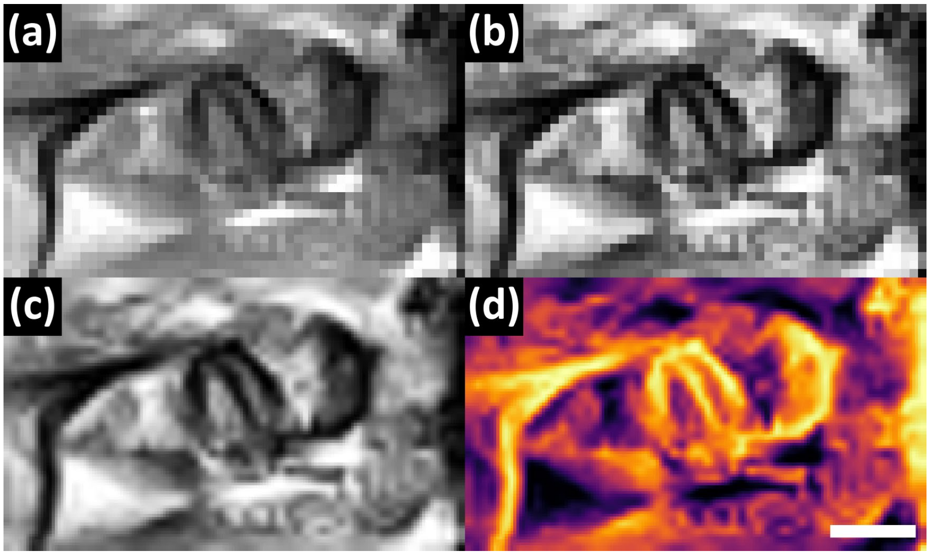


**Fig. S9: MRI image processing steps** All images in the analyzed feeding sequences were (a) cropped, (b) locally contrast-enhanced, (c) upsampled by 3x, and (d) false colored. All frames were processed with the same settings. Scale bar = 10mm. Same scale for all panels.

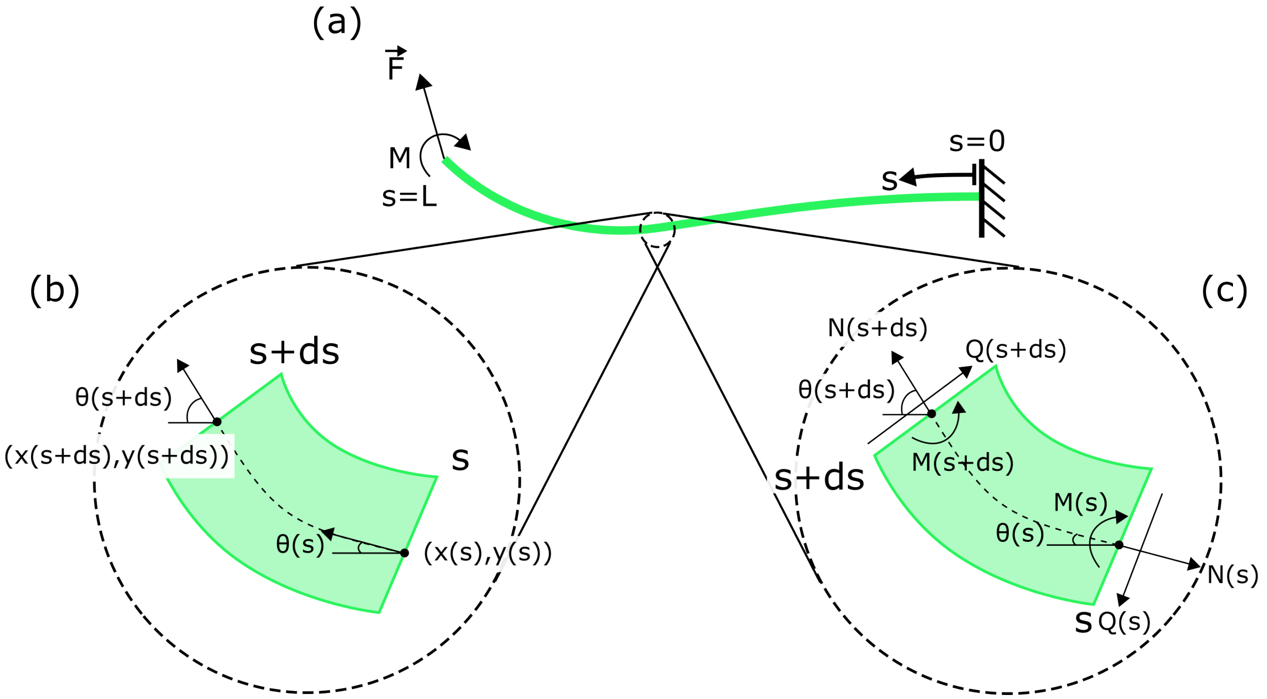


**Fig. S10: Kinematics and free body diagram of a geometrically exact Euler-Bernoulli beam** (a) A beam is a 1-dimensional continuum element capable of resisting both forces and moments. The beam geometry is parameterized by the arc length of the beam $s \in [0, L]$. Here, $ds$ is an infinitesimal change in the arc length position. (b) For each infinitesimal segment of the beam, we track the planar position $(x(s), y(s))$ and the tangent heading angle $\theta(s)$. (c) Each segment of the beam is subject to axial forces ($N(s)$), shear forces ($Q(s)$), and moments ($M(s)$). The governing equations of the beam are derived using a balance of forces and moments on each beam segment.

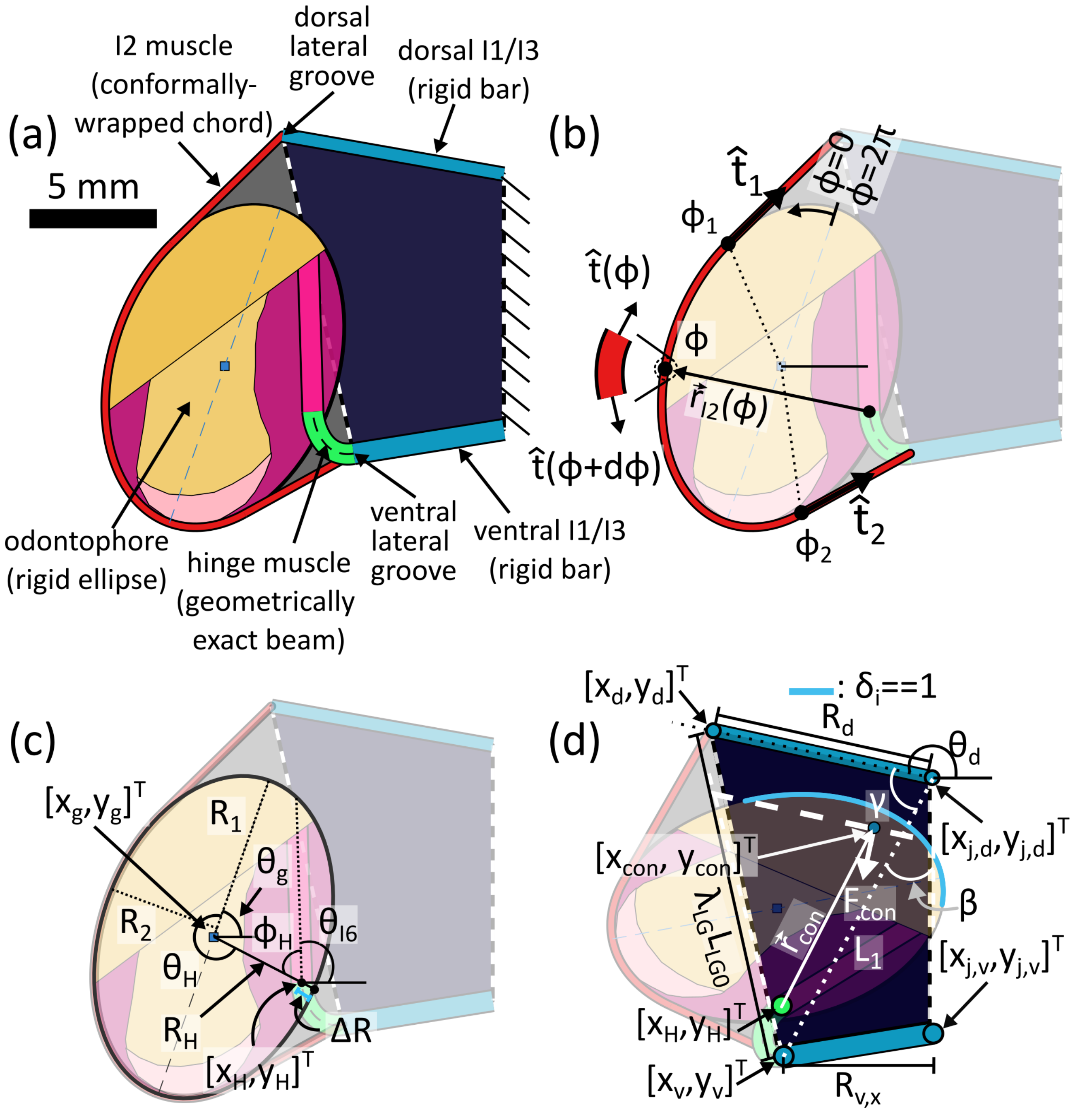


**Fig. S11: Model geometry and constraints** (a) Model geometry and component definition. (b) Geometry and mechanical advantage of the I2. The I2 wraps conformally around the odontophore ellipse from the ellipse parameter $\phi_1$ to $\phi_2$. The net force and moment are calculated by integrating the infinitesimal force vector ($\propto \partial\vec{t}/\partial\phi$) and moment ($\propto \vec{r}_{I2} \times \partial\vec{t}/\partial\phi$) between $\phi_1$ and $\phi_2$. (c) Geometry of the ellipse and hinge connection point. These geometric values are used to define the position and orientation of the odontophore ellipse for a given configuration of the hinge beam and aspect ratio of the odontophore. (d) In the model, the I3 changes shape by contracting the lateral groove and depressing the dorsal I3 line. This impacts the odontophore through a contact force and associated moment. See main manuscript text for variables definitions. The moment arms $\vec{r}_{I2}$ (for the I2 muscle) and $\vec{r}_{con}$ (for the dorsal I3 contact force) enhance the abilities of these muscles to bend the hinge structure.

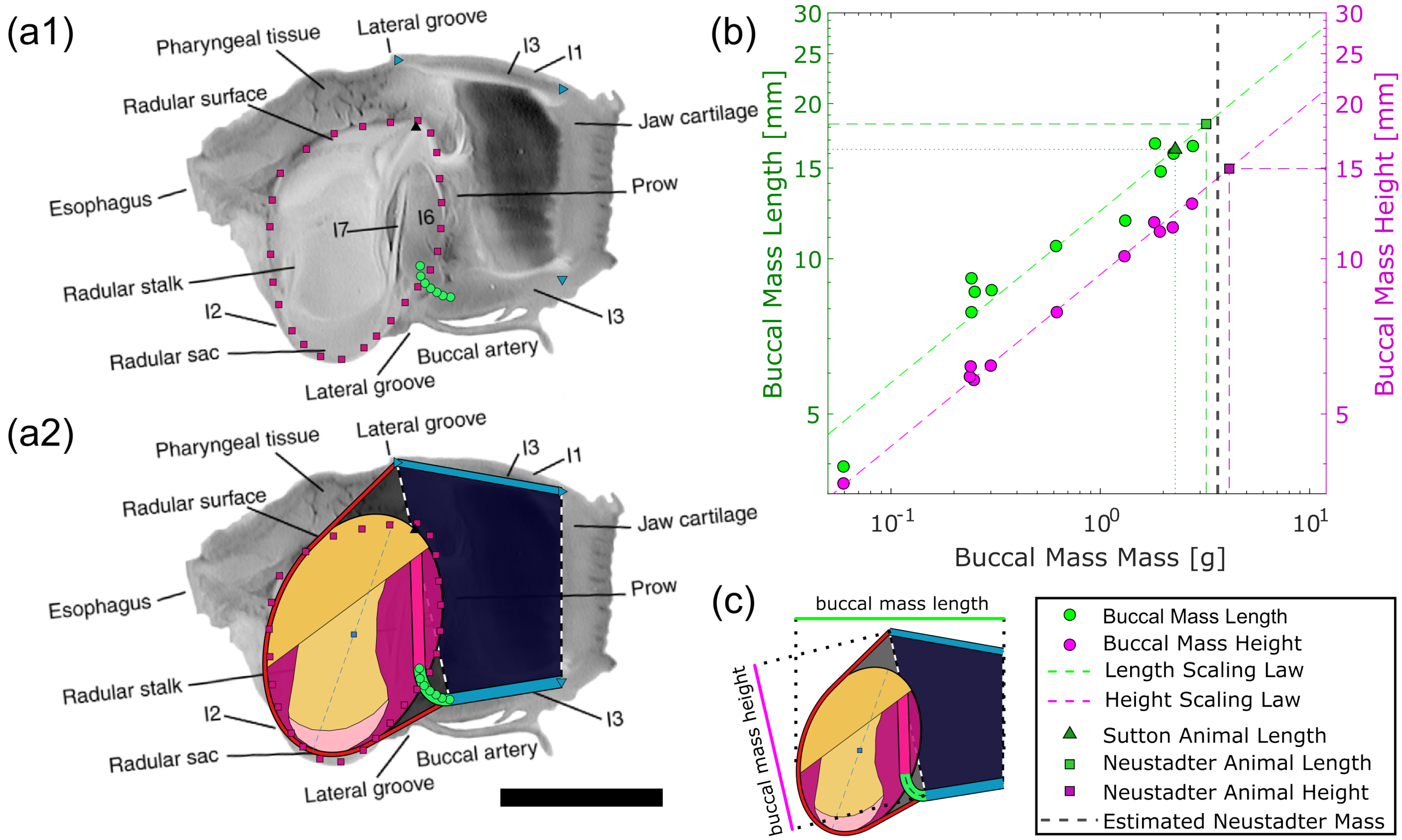


**Fig. S12: Estimating model anatomy from midsagittal and scaling data.** (a1) Data points were digitized from a lateral view of a midsagittally bisected buccal mass from Neustadter et al. 2007[S1] (modified with permission). Magenta squares: odontophore outline. Green circles: hinge midline. Blue right-facing triangle: dorsal I1. Blue down triangle: ventral I1. Black up triangle: I6. (a2) These data points were used to fit the resting geometry of the biomechanical model. The anatomical image in (a1) and (a2) is rotated to align the animal data with the biomechanical model where the jaw line is oriented vertically. Scale bar: 10mm, same for (a1) and (a2). (b) Scaling data for the buccal mass length (green) and height (magenta) as a function of mass of the buccal mass (digitized from Rogers *et al.* 2024[S4]). Isometric scaling laws were fitted to the data and used to estimate the masses of buccal masses from which we used data for model calibration. Geometric measurements for buccal masses used in Sutton *et al.* 2004[S5] (green up triangle: buccal mass length) and Neustadter *et al.* 2007[S1] (green square: buccal mass length, magenta square: buccal mass height) were used to estimate the masses of the buccal mass for those data sets. (c) Buccal mass length is defined as the largest anteroposterior length of the buccal mass, and the height of the buccal mass is defined as the length of the lateral groove (definitions based on Rogers *et al.* 2024[S4]).
